# Unlocking viral evasion: Luminal charge interactions in BoHV-1 UL49.5 allosterically control TAP degradation

**DOI:** 10.64898/2026.06.11.731699

**Authors:** Natalia Karska, Małgorzata Graul, Igor Zhukov, Sylwia Rodziewicz-Motowidło, Andrea D. Lipińska, Magdalena J. Ślusarz

## Abstract

The UL49.5 protein of bovine alphaherpesvirus 1 (BoHV-1) is known to inhibit the transporter associated with antigen processing (TAP) and interfere with antigen presentation, in part by promoting TAP degradation. However, the role of electrostatic interactions within the N-terminal luminal domain in controlling these processes remains unclear. Here, we combined circular dichroism (CD), solution nuclear magnetic resonance spectroscopy (NMR), all-atom molecular dynamics simulations, and cell-based assays to define the structural and functional contribution of charged residues within the *N*-terminal luminal domain of UL49.5. Two *N*-terminal variants of UL49.5, UL49.5^22-56^RR(30–31)DD and UL49.5^22-56^D36K, with substitutions of charged-reversal amino acid residues, were designed. These two mutants formed membrane-induced α-helical structures but showed altered helix stability and interaction patterns. Molecular dynamics simulations of the UL49.5–TAP complexes revealed that wild-type UL49.5 forms a stable electrostatic interface with TAP, particularly in the unkinked conformation, while charge-reversal mutations remodel salt-bridge networks, destabilize the luminal helix, and alter the positioning and dynamics of the transmembrane and cytoplasmic *C*-terminal regions.

The structural changes within the *N*-terminus alter the exposure of the *C*-terminal degron required for KLHDC3-dependent degradation. Consistent with these findings, the mutants did not induce proteasomal degradation of TAP, despite maintaining near wild-type levels of downregulation of MHC class I. Together, these results identify *N*-terminal electrostatic interactions as allosteric determinants of UL49.5-driven TAP degradation and demonstrate that TAP degradation can be mechanically uncoupled from downregulation of MHC class I. This study improves our understanding of viral immune evasion strategies and potential therapeutic targets.

## 1. Introduction

Bovine alphaherpesvirus 1 (BoHV-1) is a significant pathogen within the *Alphaherpesvirinae* subfamily and the genus *Varicellovirus*, known for its detrimental effects on cattle health. BoHV-1 is responsible for causing a variety of diseases in cattle, including conjunctivitis, respiratory tract infections, abortion, encephalitis, and fatal systemic infections (1). The BoHV-1 genome encodes a multitude of proteins, approximately 70 to 80, that are involved in viral replication, assembly, latency, and evasion of host immune responses (2).

Among these proteins, UL49.5 has emerged as a critical immune evasion factor, playing a pivotal role in inhibiting the Transporter Associated with Antigen Processing (TAP) of diverse mammalian origin, including human TAP, and reducing the expression of class I molecules of the major histocompatibility complex (MHC) on the surface of host cells (3). UL49.5 is a type 1 transmembrane protein consisting of 96 amino acid residues, with a predicted molecular mass of approximately 9 kDa (4). It is composed of a cleavable *N*-terminal signal peptide, a non-glycosylated ER-luminal domain, a transmembrane helix, and a *C*-terminal cytosolic tail, each playing a specific role in the protein’s function (4,5).

The luminal domain of UL49.5 is essential for inhibiting TAP and contains a conserved cysteine residue that is believed to stabilize the protein via disulfide bonding with virus glycoprotein M (gM), thus interfering with peptide translocation by TAP (4,6). The membrane-proximal *N*-terminal residues and the transmembrane region anchor UL49.5 in the endoplasmic reticulum (ER) membrane, mediating interactions with TAP subunits, and contributing to the non-functional conformation of TAP (7). The cytoplasmic tail, while not essential for TAP inhibition, plays a crucial role in inducing TAP degradation through the ER-associated degradation pathway (ERAD) (3).

Recent cryo-EM analyses have resolved the overall architecture of the BoHV-1 UL49.5–TAP inhibitory complex, showing that UL49.5 occupies the TAP translocation pathway and engages TAP through its luminal and transmembrane regions. However, the detailed conformation of the *N*-terminal luminal segment within the TAP cavity remains poorly resolved. This leaves unclear how local electrostatic interactions in this region contribute to the stability of the UL49.5–TAP complex, TAP inhibition, and downstream TAP degradation (8). Recent studies have shed light on the complex interplay of multiple motifs within UL49.5, including the RRE (30–32) motif and an unstructured fragment PPQ (52–54) forming a ‘proline hinge’ (Graul et al., 2023; Karska et al., 2019, 2021). Structural analyzes have highlighted the importance of the RRE motif in stabilizing the α-helical structure of the luminal domain UL49.5, with mutations in this motif affecting protein flexibility and TAP inhibition (11). We could previously identify the PPQ motif as crucial for UL49.5’s ability to inhibit TAP, with mutations in this region that impact the expression of class I MHC and TAP levels (7). Moreover, the PPQ motif was found important for the rearrangement of the UL49.5 helices upon binding to the kelch domain-containing protein 3 (KLHDC3) - a substrate receptor of a cullin 2-RING (CRL2) E3 ubiquitin ligase complex of the DesCEND (destruction via a *C*-end degrons) pathway (8). The primary objective of this study was to investigate the role of the UL49.5 luminal domain in TAP inhibition and proteasomal degradation by introducing targeted point mutations that alter the charge within the *N*-terminal luminal region of UL49.5. Through a combination of circular dichroism (CD), nuclear magnetic resonance (NMR) spectroscopy, molecular modeling in an endoplasmic reticulum-mimetic membrane environment, and cellular functional assays, we aimed to elucidate the influence of specific electrostatic determinants within the luminal domain on UL49.5 conformation, its interaction with TAP, and the subsequent effects on TAP stability and MHC class I surface expression.

In this study, we hypothesized that altering the charge distribution within the luminal domain of UL49.5 would affect its ability to inhibit TAP and modulate MHC class I expression. However, although the *C*-terminal RGRG degron of UL49.5 has been implicated in KLHDC3-dependent TAP degradation, it remains unclear how contacts formed by the ER-luminal *N*-terminal region are coupled to the presentation of the degron on the cytoplasmic side of the membrane. Therefore, we tested whether charge-reversal mutations within the luminal helix alter the stability of the UL49.5–TAP interface, the accessibility of the *C-*terminal degron and the functional relationship between TAP degradation and MHC class I downregulation.

## 2. Results

### Structural properties of the designed peptides

To elucidate the structural contribution of the endoplasmic domain, we designed two peptide analogs based on the native UL49.5 sequence (Figure 1). We used these to assess how specific modifications in charge distribution influence its conformation and potential interaction with TAP. In the first analog, UL49.5^22-56^RR(30–31)DD, we substituted two positively charged arginine residues at positions 30 and 31 with negatively charged aspartic acids, resulting in a reversal of the local electrostatic potential. In the second analog, UL49.5^22-56^D36K, a negatively charged aspartic acid residue was substituted with lysine, thus engendering a local positive charge at the interface interacting with TAP. Substitutions of amino acid residues in the luminal domain were aimed at changing the local electrostatic surface, which could potentially affect the conformational stability of the peptide and consequently the affinity of the UL49.5 mutants for TAP. Analogs in which the polarization of charge changes at specific positions provide a basis for investigating the effect of electrostatic complementarity and local charge shifts on the mechanism of interaction between UL49.5 and TAP. To avoid problems related to dimerization by oxidation of the cysteine sulfhydryl group and/or formation of diastereomeric sulfoxides by oxidation of the thioether moiety in methionine, methionine and cysteine were replaced by isosteric norleucine (Nle) and (S)-2-aminobutyric acid (Abu), respectively.

**Figure 1.**
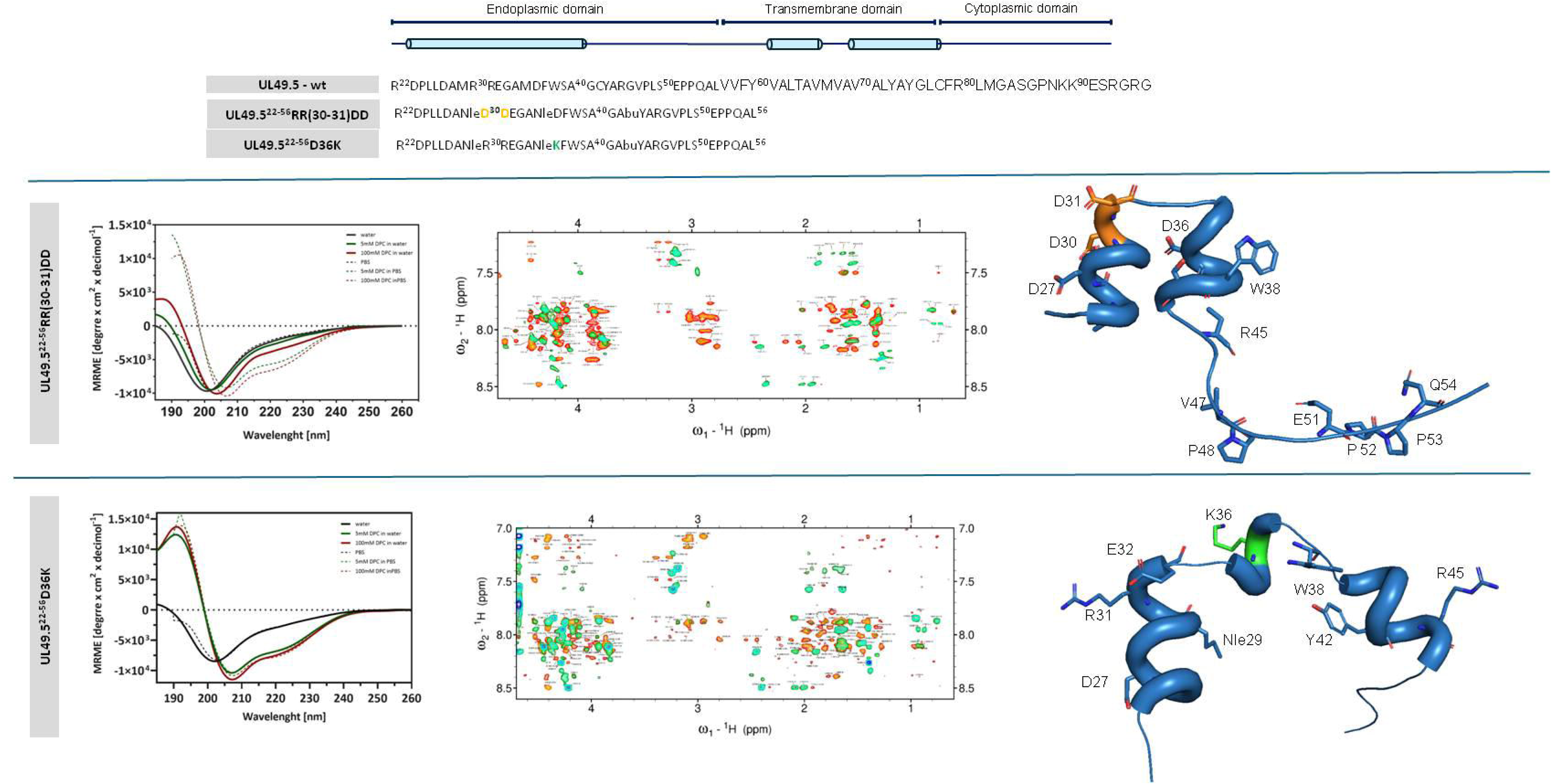
Top panel: A comparison of the sequence of the native UL49.5 N-terminal fragment and its analogs. Modified amino acid residues have been marked in orange and green. Bottom panel: Structural characterization of the peptides UL49.5^22-56^RR(30–31)DD and UL49.5^22-56^D36K derived from BoHV-1. The upper row presents biophysical and structural data for the UL49.522-56RR(30–31)DD peptide. The bottom row shows analogous data for the UL49.5^22-56^D36K peptide. CD spectra indicate that both peptides display α-helical characteristics in a micellar environment, whereas in water and PBS they exhibit features of a disordered conformation. The overlaid NMR spectra show TOCSY spectra (green) superimposed on the NOEY spectra (orange), revealing the presence of medium-range Nuclear Overhauser effect (NOE) contacts, which are characteristic of α-helical segments. The right-hand panels show three-dimensional NMR-derived structures of mutant peptides. Two helical regions are observed that span residues P24–D31 and Nle35-S39 for the UL49.5^22-56^RR(30–31)DD peptide. By contrast, the UL49.5^22**-**56^D36K peptide contains three helical regions located at P24–E32, A34–F37 and A40–G46.

### Revealing the structure–charge relationship in UL49.5 peptide analogs

To investigate how changes in the structure of the luminal UL49.5 domain affect its interaction with TAP, two designed peptides, UL49.5^22-56^RR(30–31)DD and UL49.5^22-56^D36K, were studied using CD and solution NMR spectroscopy. This experimental strategy was based on our previous studies showing that the structural organization of UL49.5-derived fragments is strongly dependent on membrane-mimetic conditions (10). These complementary techniques were used to understand how the luminal segment responds structurally to membrane-mimetic conditions and sequence-specific charge modifications introduced within this domain.

CD spectroscopy was utilized to identify changes in the secondary structure of the peptides in the presence of various membrane environments at neutral pH. The results revealed clear environment-dependent changes in the secondary structure of the peptides. In the absence of DPC micelles, the peptides exhibited an unordered structure, as indicated by the minima in the mean residue molar ellipticity (MRME) near a wavelength of 199 nm and positive values at a wavelength of around 220 nm (Figure 1). However, in the presence of detergent micelles, the peptides adopted a mainly α-helical structure, as indicated by MRME minima near 208 and 222 nm.

The conformational properties of the mutant peptides derived from the luminal domain of the UL49.5 protein in a membrane-mimetic environment were further characterized using solution NMR spectroscopy. Complete resonance assignments of all residues for the ^1^H, ^15^N and 13C nuclei were obtained for both peptides in DPC-d₃₈ micelles using a combination of two-dimensional homonuclear TOCSY and NOESY experiments, complemented by heteronuclear 1H–^13^C and ^1^H–^15^N HSQC spectra (Figure 1S-5S). Analysis of NOESY spectra revealed the characteristic medium-range NOE cross-peaks of the α-helical conformations, consistent with the secondary structure inferred from CD measurements under micellar conditions.

A quantitative analysis of the NOE-derived distance restraints revealed distinct distributions for the two mutant peptides, indicating differences in the extent and continuity of their ordered secondary structures. The NMR-derived structures of the mutant peptides showed that both peptides displayed α-helical characteristics in a micellar environment. The NMR-derived structures of the mutant peptides showed two helical regions spanning residues P24–D31 and Nle35–S39 for the UL49.5^22-56^RR(30–31)DD peptide, while three helical regions were located in P24-E32, A34-F37 and A40–G46 for the UL49.5^22-56^D.

Structure calculations were performed using NOE distance restraints derived from experiments, as well as hydrogen bond and backbone torsion angle restraints. The Ramachandran plot statistics showed that a high percentage of residues were located in the regions most favored for both peptides, with acceptable RMS Z-scores for bond lengths, bond angles, dihedral angles and non-bonded interactions, supporting the stereochemical consistency of the calculated structures (Table 1S). The three-dimensional structures derived from NMR revealed different helical architectures for the two mutant peptides, with the UL49.5^22-56^D36K peptide showing a more segmented helical organization compared to the UL49.5^22-56^RR(30–31)DD peptide.

**Table 1.** Peptide purification and identification.

| Peptide | Theoretical mass | Observed mass | Purification conditions |
| --- | --- | --- | --- |
| UL49.5<sup>22-56</sup>RR(30-31)DD | 3742,72 | 3742,7 | 35% – 50% B in 90 min |
| UL49.5<sup>22-56</sup>D36K | 3838 | 3838,1 | 40% – 80% B in 90 min |

### Structural organization and dynamics of UL49.5–TAP complexes in the membrane

The structural organization and dynamics of the UL49.5–TAP complexes in the membrane were investigated using 1 μs all-atom molecular dynamics (MD) of the wild-type UL49.5 protein and its mutants (UL49.5-RR(30–31)DD and UL49.5-D36K) in two different conformational states: unkinked and kinked. These two conformations represent alternative structural states of the transporter, the unkinked form dominant at 56%, whereas the kinked state represents 27% (12). The structural properties of UL49.5 variants were analyzed separately for complexes with TAP in the unkinked and kinked conformations.

In complexes with TAP in an unkinked conformation, the wild-type UL49.5 protein formed an α-helical segment in the luminal region spanning residues A28–W38, with a well-defined transmembrane helix from F60 to R80 (Figure 7S). The largest fluctuations were observed in the cytoplasmic *C*-terminal segment (Figure 8S). The mutant UL49.5-RR(30–31) DD exhibited an overall structural organization similar to the wild-type protein (Figure 2), with a reduced α-helical content in the luminal region and increased flexibility in the cytoplasmic *C*-terminal region (Figure 8S). The mutant UL49.5-D36K displayed a partially helical organization in the luminal region, with fragmented helical segments (Figure 7S) and increased flexibility in the cytoplasmic *C*-terminal region (Figure 8S).

**Figure 2.**
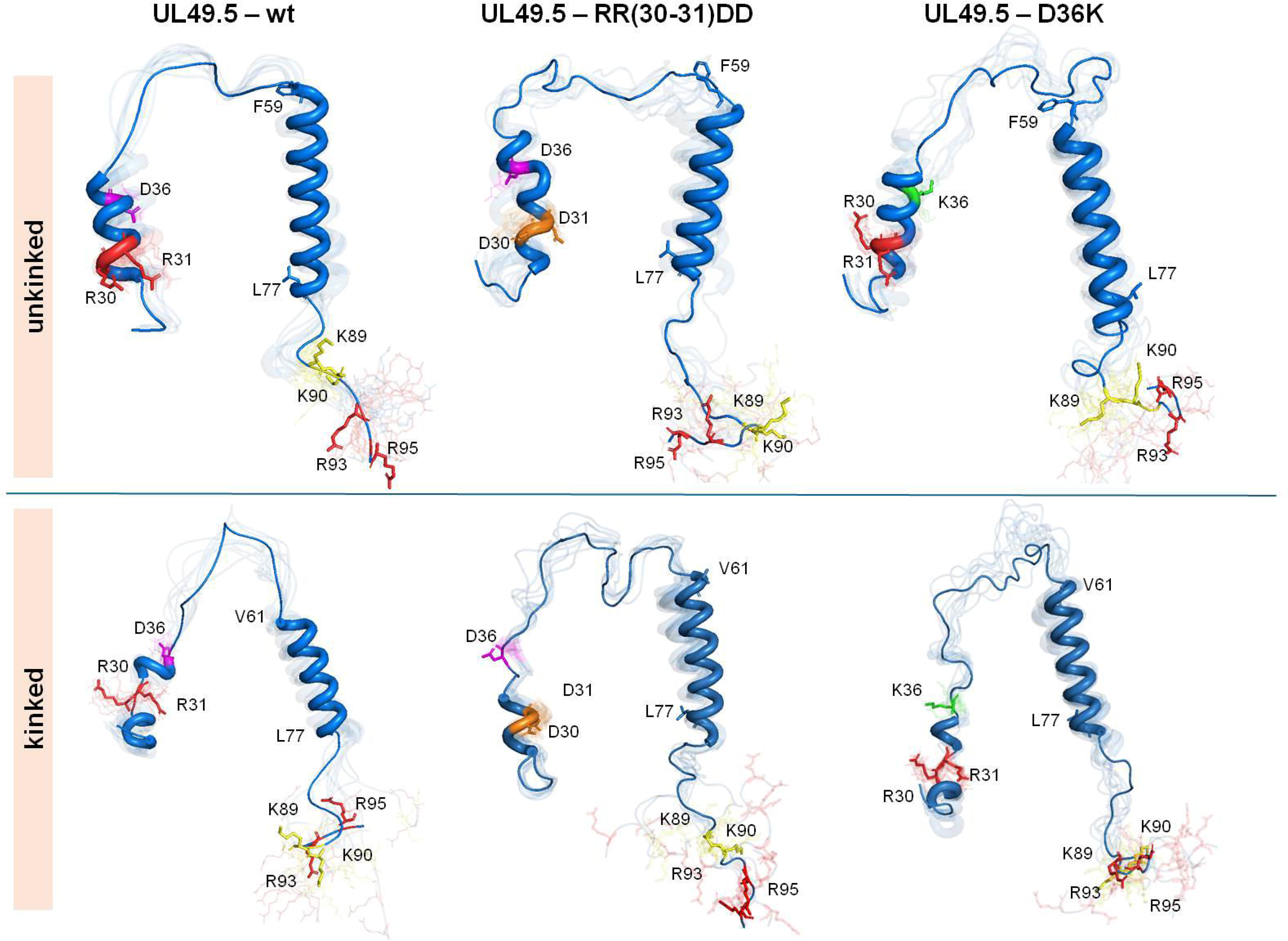
Structures of UL49.5 variants after molecular dynamics simulations. Representative structures of the wild-type UL49.5 protein and two mutants (UL49.5-RR(30–31)DD and UL49.5-D36K) obtained after molecular dynamics simulations. These structures are presented for complexes formed with TAP in two distinct conformations: unkinked (top row) and kinked (bottom row). Selected residues in the N-terminal (R30, R31 and D36) and C-terminal (K89, K90, R93 and R95) regions are highlighted to indicate the positions of mutations and degron residues relevant to interactions with KLHDC3.

In the kinked TAP conformation, the wild-type UL49.5 protein adopted a stable helical architecture with two short α-helical segments in the luminal region and a transmembrane helix spanning residues F59–L77 (Figure 2). The transmembrane helix remained predominantly α-helical, with minor fluctuations at the termini, while the luminal region showed increased structural variability, particularly around residues R30 and R31. The *C*-terminal segment of the cytoplasm exhibited relatively low fluctuations, with increased mobility observed at the terminal residues (Figure 8S). The mutant UL49.5-RR(30–31)DD showed a different structural organization, forming a continuous α-helical segment in the luminal region spanning residues D27–W38, with a conserved transmembrane helix from V61 to C78 (Figure 2). Mutations at positions 30 and 31 resulted in increased structural heterogeneity within the luminal segment (Figure 7S), with higher values of RMSF observed in the cytoplasmic *C*-terminal region (Figure 8S). The mutant UL49.5-D36K, with a charge-reversal substitution at position 36, exhibited a continuous α-helical segment in the luminal region spanning residues A28–S39, while maintaining a transmembrane helix from Y60 to F79 (Figure 2). The *C*-terminal segment of the cytoplasm remained largely disordered, with increased fluctuations observed in the *C*-terminal region (Figure 8S).

In general, MD simulations provided insight into the structural organization and dynamics of UL49.5–TAP complexes. It should be noted that among all complexes analyzed, the highest content of *N*-terminal helix was observed in the wild-type UL49.5 complex with unkinked TAP, representing the major cryo-EM population (Figure 7S). Moreover, in this complex, the *C*-terminal segment of UL49.5 was the most stable among all investigated systems with the lowest RMSF values (Figure 8S).

### Interaction interface between UL49.5 and TAP

The interactions between UL49.5 and the TAP transporter were analyzed to understand the mechanism of viral inhibition of peptide transport. The analysis of MD trajectories revealed that the luminal domain of UL49.5 inserts into the transmembrane cavity of TAP and interacts with residues in its luminal region. The interaction interface primarily involves residues within the *N*-terminal luminal region of UL49.5 (Figure 9S). Analysis was conducted for the wild-type UL49.5 protein and two mutants (UL49.5-RR(30–31)DD and UL49.5-D36K) in complexes with TAP in unkinked and kinked conformations.

In the unkinked conformation of TAP, R22 of UL49.5 forms salt bridges with E201, E242, and D246 of the TAP1 subunit, while R30 interacts with E242 (TAP1) and E417 (TAP2), and forms a hydrogen bond with R381 (TAP2). E32 interacts with R273 and K277 of TAP2, and D36 forms salt bridges with R312 and hydrogen bonds with Q453 and Y250 of TAP1, positioning the luminal region of UL49.5 at the luminal opening of the TAP cavity (Figure 3 and 9S).

**Figure 3.**
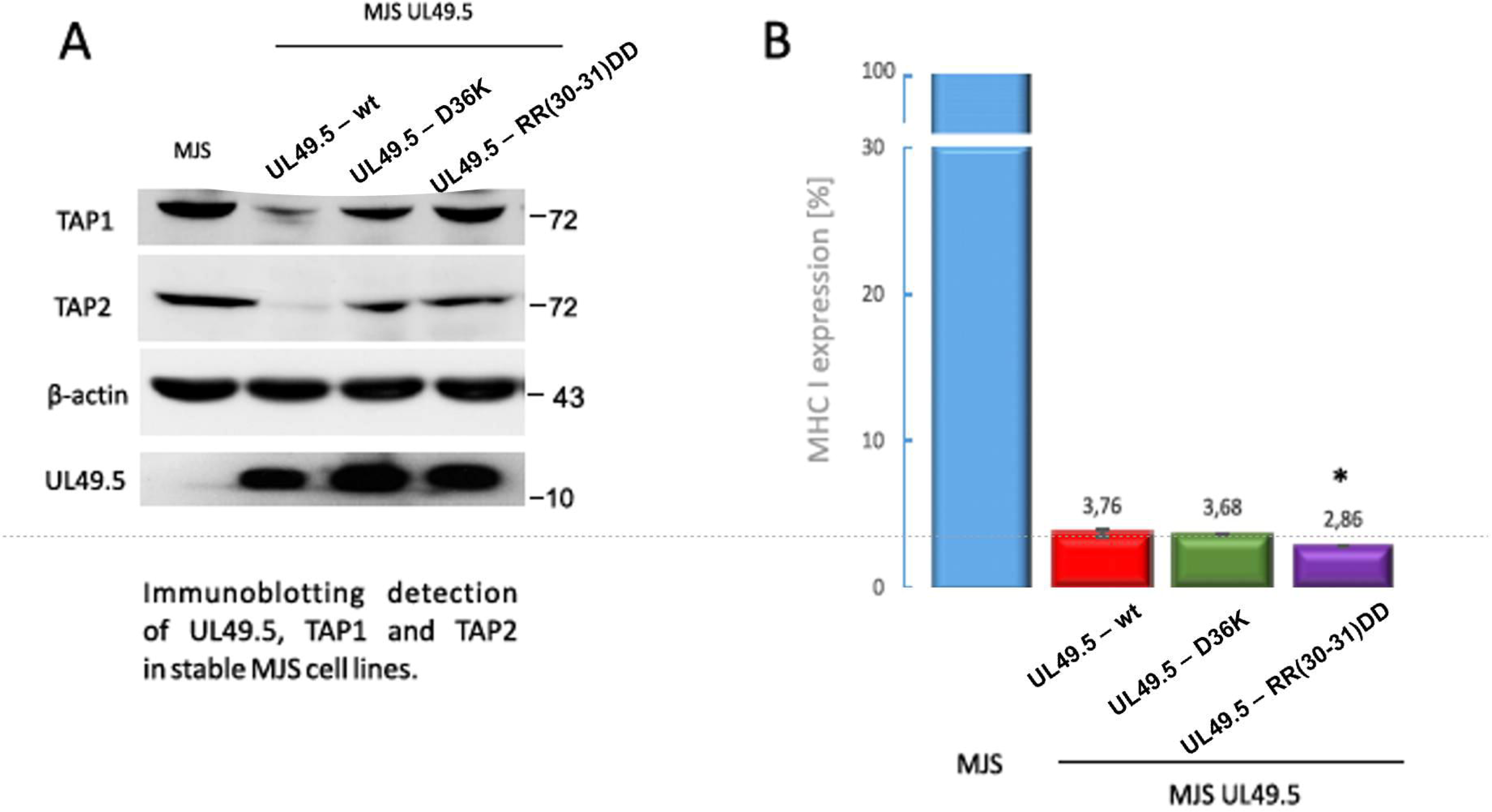
Interactions between UL49.5 (wt and mutant variants RR(30–31)DD and D36K) and TAP observed in unkinked (top) and kinked (bottom) conformations during 1 µs MD simulations. Representative conformations illustrate the spatial arrangement of UL49.5 relative to the TAP complex in both states. TAP1 is shown in yellow, whereas TAP2 is shown in green.

In the UL49.5-RR(30–31)DD mutant, the substitution of R30 and R31 with negatively charged residues reorganizes the interaction interface. Acid residues within the luminal region mediate electrostatic contacts, with D27, D30, and D31 forming salt bridges with R273 and K277 of the TAP2 subunit. Contacts with the TAP1 subunit are partly preserved through residues R22 and E32, although the interaction mediated by E32 shifts towards TAP1 in the mutant.

In the UL49.5-D36K mutant, the interactions are primarily mediated by residues in the N-terminal segment, with R22 forming salt bridges with D246 and E242 of TAP1. Additional contacts involve D23, which interacts with R373 of TAP2, and R30, which forms a salt bridge with E242 of TAP1 and hydrogen bonds with S238 and E201 of TAP1.

In complexes with kinked TAP, the polar interactions are established with residues from both TAP subunits, with R22 forming salt bridges with E417 and E242, and D23 interacting with R380 and E201. R30 forms stabilizing contacts through the salt bridges with E242 and E201 of TAP1 and E417 of TAP2, while E32 forms a salt bridge with R312, and D36 forms a hydrogen bond with K277 and E453.

In the UL49.5-RR(30–31)DD mutant, the reorganization of the interaction interface observed for UL49.5-wt leads to preserved contacts with TAP2 and new contacts within the luminal region. D23 interacts with R373, and D31 forms a salt bridge with K277, while E32 interacts with both TAP subunits. In the complex of the UL49.5-D36K mutant with kinked TAP, the substitution of D36 with lysine leads to a partial reorganization of the interaction within the luminal region, with R22 forming a salt bridge with E242.and R30 interacting with E417 and E242. Additional contacts involve D23, D27, and R31 with various residues of the TAP subunits.

In general, molecular dynamics simulations provided information on the interaction interface between UL49.5 and TAP in unkinked and kinked conformations, highlighting the role of specific residues in mediating these interactions. The highest number of strong salt bridges was observed in the wild-type UL49.5 complex with unkinked TAP (see Figure 9S). These salt bridges are evenly distributed throughout the UL49.5 structure, engaging both the *N*- and *C*-termini, resulting in enhanced overall stability of this complex. This was confirmed by the linear interaction energy (LIE) values (Table 2S). The average LIE value for the wild-type UL49.5 complex with unkinked TAP was approximately −1077 kcal/mol, which was the lowest among all investigated systems. Any of the introduced mutations weakens the interaction between the proteins, resulting in less stable complexes with average LIE values of −807 kcal/mol and −733 kcal / mol for the UL49.5-RR(30–31)DD and UL49.5-D36K complexes, respectively. In contrast, for the wild-type UL49.5 complex with kinked TAP, which is much less stable (average LIE = −846 kcal/mol), the effects of introduced mutations vary: the RR(30–31)DD mutation improves protein-protein interaction and stability of the complex (average LIE = −988 kcal / mol), while the D36K mutation results in a less stable complex (average LIE = −814 kcal / mol).). The LIE values confirm the highest stability of the wild-type UL49.5-TAP complex, with the UL49.5 structure characterized by a stable *N*-terminal helix and a moderately mobile *C*-terminus, as described in the previous section.

**Table 2.**
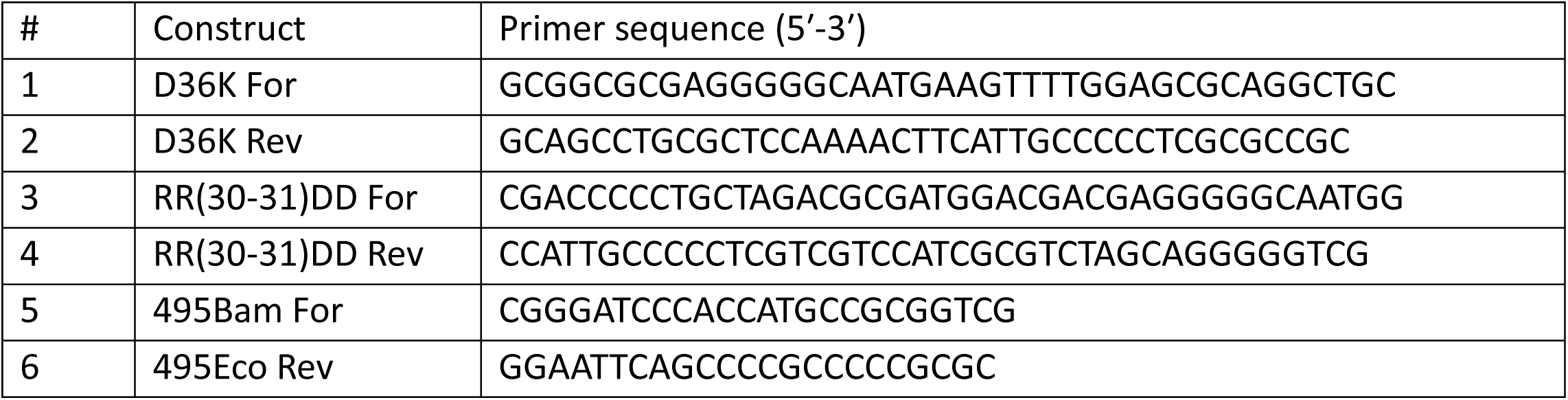
List of PCR primers used in the study. Primers #1–4 were used for site-directed mutagenesis of UL49.5. For: forward; Rev: reverse.

### Structural dynamics of the UL49.5 *C*-terminus and exposure of the degron

The cytoplasmic *C*-terminal region of UL49.5 was analyzed to investigate the conformational dynamics and solvent exposure of the *C*-terminal degron RGRG (93–96), which is responsible for UL49.5 presentation to the cellular degradation machinery (8). The analysis focused on the degron and the preceding UL49.5 residues 87–92. In the wild-type UL49.5 protein, it was found that the *C*-terminal region is solvent-exposed in both TAP conformations, indicating a flexible cytoplasmic tail (Figure 3 and Figure 10S). However, the exposure of this region was modulated by the conformational state of the transporter.

In the kinked conformation of TAP, the solvent accessibility of the segment 87 to 96 was higher than in complexes of the unkinked TAP conformation, suggesting high conformational mobility and exposure to degron. In the unkinked state of TAP, in the complexes with mutants, slightly reduced solvent accessibility was observed (Figure 10S) and, more importantly, the lowest average SASA value, 849.52 Å², was recorded for the *C*-terminus of the wild-type UL49.5 complexed with unkinked TAP. This differs significantly from the values obtained for the other systems. In addition, to assess changes in the position of the TM helix adjacent to the *C*-terminus, we analyzed the distance between the termini of the UL49.5 helices (P24-F79, Figure 11S). During the MD simulations, the helices could change their positions relative to each other due to conformational changes within the connecting loop containing the proline hinge motif. Because the *N*-terminal helix was tightly bound within the TAP pocket, changes in the calculated distance allowed us to assess how the TM helix is tilted relative to the TAP. The most significant fluctuations throughout the MD simulation were observed for both complexes of wild-type UL49.5. However, after 800 ns, the position of the TM helix stabilized, maintaining a distance of P24-F79 of approximately 37 Å (Figure 11S) with the *C*-terminal fragment of the TM helix noticeably displaced from the TAP (Figure 3). In mutant complexes, the *C*-terminus of the TM helix remained closer to the TAP, with the distance of P24-F79 ranging from 29 to 33 Å throughout MD simulations (Figure 3 and 11S).

These results suggest that the accessibility of the RGRG degron is controlled by conformational coupling between UL49.5 and TAP. Mutations within the *N*-terminal luminal helix of UL49.5 can modulate solvent exposure and mobility of the *C*-terminal segment, as well as the position of the TM helix. Beyond solvent accessibility, the orientation of the *C*-terminus is likely critical for degron recognition. In the complex of UL49.5 with unkinked TAP, the *C*-terminus showed limited but stable exposure with the position of the transmembrane helix displaced from the TAP, probably being a productive conformation of degron. In the remaining complexes, the *C*-terminal part of the transmembrane helix is located closer to TAP, which may prevent KLHDC3 from recognizing the degron, despite increased flexibility or solvent exposure of the *C*-terminus.

### Biological impact of UL49.5-D36K and UL49.5-RR(30–31)DD mutations on antigen presentation and TAP degradation

To investigate the role of the *N*-terminal electrostatic interactions of UL49.5 in TAP degradation and MHC class I downregulation, two gene constructs encoding the *N*-terminal mutants, UL49.5-D36K and UL49.5-RR(30–31)DD, were produced and delivered to MJS cells using retrovirus vectors to obtain stable cell lines, as previously described (7). UL49.5 protein levels were comparable across the generated cell lines, as demonstrated by immunoblotting (Figure 4A). In the same cell lysates, TAP1 and TAP2 levels were significantly reduced in cells expressing UL49.5-wt but not in cells expressing UL49.5-D36K or UL49.5-RR(30–31)DD. This indicates that both mutations prevent the degradation of TAP driven by UL49.5.

**Figure 4.**
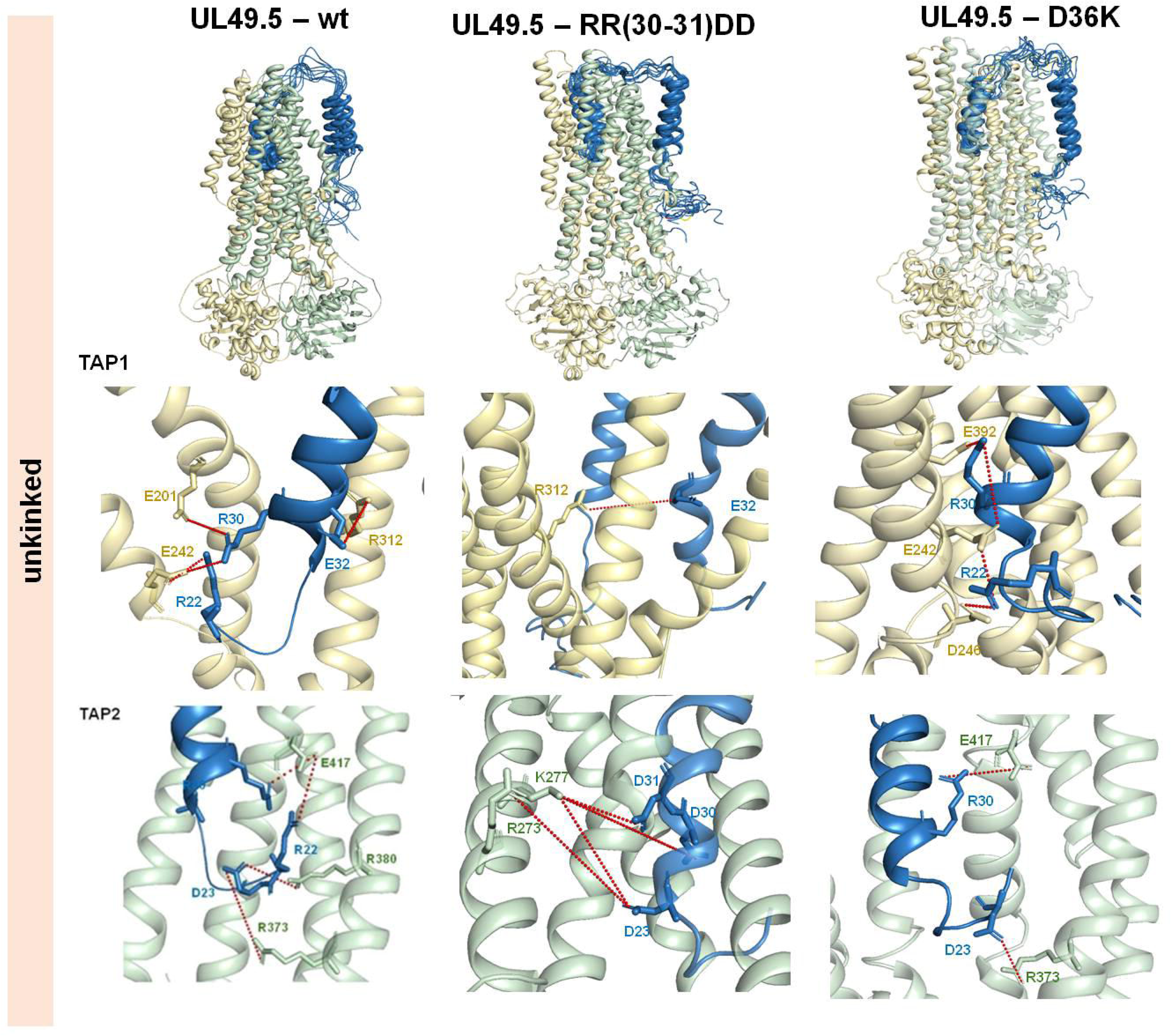
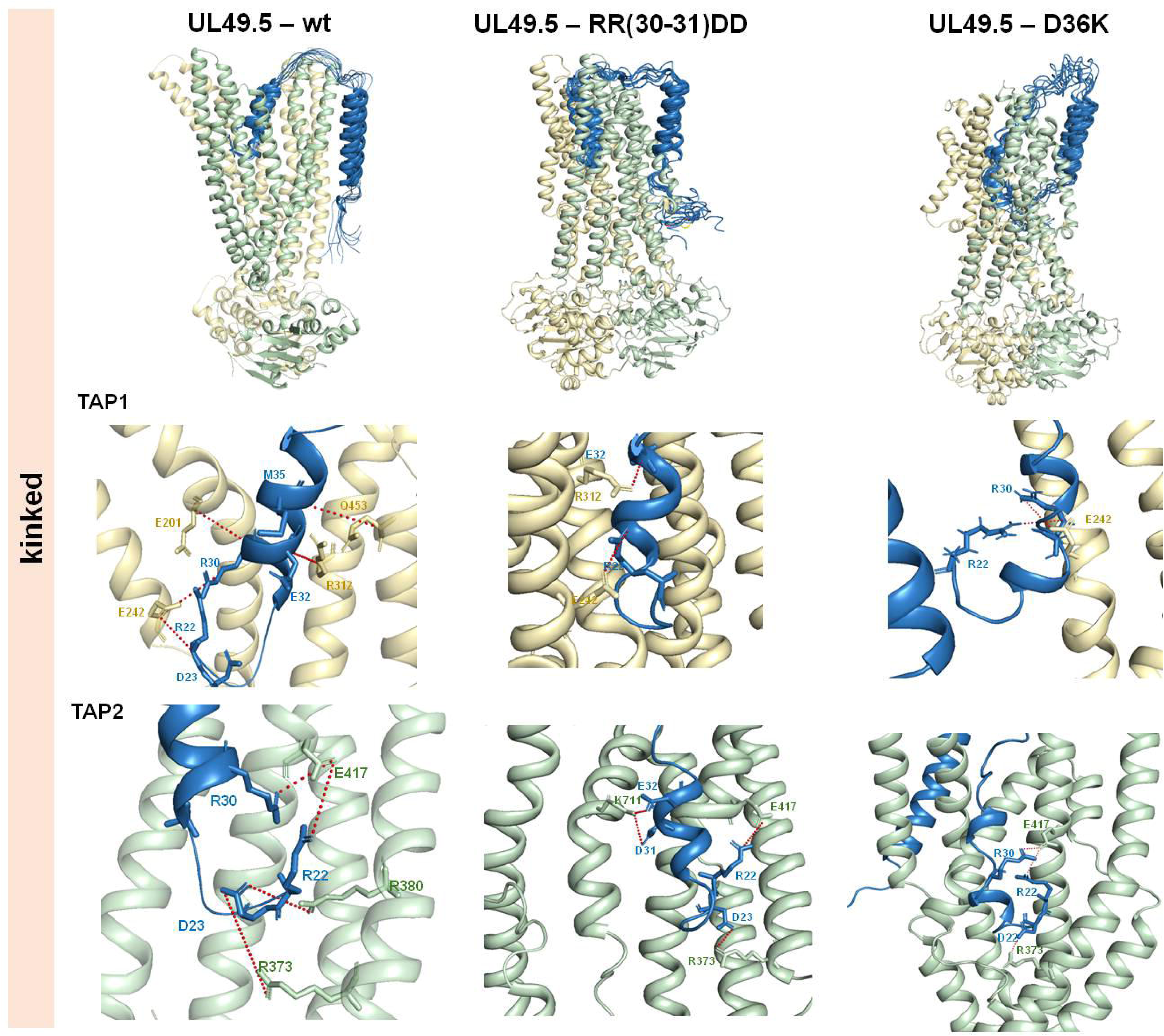
N-terminal charge-reversal mutations abolish UL49.5-mediated TAP degradation but not MHC class I downregulation. A) The expression of UL49.5 variants and the levels of TAP1 and TAP2 subunits in stable MJS cell lines was determined by immunoblotting using specific antibodies; β-actin was used as a loading control. B) The cell surface expression of MHC class I in MJS cells was evaluated by flow cytometry using specific antibodies (W6/32); and presented as the percentage of mean fluorescence intensity; the fluorescence of parental cells without UL49.5 was set as 100%. The analysis was performed in triplicate. The statistical significance of differences between MJS UL49.5wt and MJS with UL49.5 variants was estimated by one-way ANOVA;; *p<0,2.

The cell surface expression of MHC class I molecules on MJS cell lines was determined by flow cytometry. Compared to the control cells (MJS), UL49.5 WT reduced MHC class I surface levels to a small fraction of the control. D36K and RR(30–31)DD mutations in UL49.5 did not significantly affect efficiency in MHC class I downregulation (Figure 4B). Together, these data demonstrate that the *N*-terminal salt bridges in UL49.5 may regulate the degradation of TAP.

## 3. Discussion

Recent cryo-electron microscopy studies have provided valuable insight into the structural basis of the inhibitory complex formed between the BoHV-1 UL49.5 protein and TAP (12). These studies have shed light on the interactions between UL49.5 and TAP, highlighting the role of electrostatic contacts in inhibition of peptide transport. However, the detailed organization of the *N*-terminal luminal domain of UL49.5 within the TAP cavity and the functional consequences of its electrostatic interactions could not be fully resolved by cryo-EM analysis. Our study aimed to further elucidate the structural and functional implications of specific electrostatic determinants within the luminal *N*-teriminal luminal domain of UL49.5 in the context of inhibition of TAP and downregulation of the MHC class I.

Our combined experimental and computational approach revealed that the luminal domain of UL49.5 has an intrinsic propensity to form α-helices under membrane-mimicking conditions, consistent with the cryo-EM structures showing UL49.5 inserted into the TAP translocation cavity (12). This membrane-dependent helix formation suggests that the luminal segment adopts a structured conformation when engaged with the TAP complex and the membrane environment. Importantly, our results demonstrate that modifications of the polar residues significantly impact the helical content and structural organization of the *N*-terminal helix. Mutations such as RR(30–31)DD disrupt the continuity of the helix and increase structural flexibility, while the D36K mutation preserves a more coherent helical organization, indicating the role of specific charge distributions in stabilizing the functional conformation of the luminal helix.

Our MD simulations of full-length mature UL49.5 and its mutant variants in complex with TAP within a membrane environment provided further information on the structural behavior of UL49.5. The wild-type UL49.5–TAP complex exhibited stable position within the membrane, with the transmembrane helix serving as a stable anchor and structural scaffold (12). Mutations in the luminal domain did not affect the structural integrity or stability of the transmembrane helix, highlighting its role in preserving the overall architecture of UL49.5 and ensuring the correct positioning of the inhibitory luminal domain in relation to TAP.

In contrast, mutations in the luminal domain altered the positional and structural stability of the luminal helix. The wild-type protein maintained a stable and consistent orientation relative to that of TAP, supporting the suggestion of persistent inhibitory contacts within the translocation pathway. Our simulations confirmed the structural stability of the luminal helix on microsecond timescales, with the RR(30–31)DD mutation leading to the most significant destabilizing effect, resulting in increased positional variability, helix shortening, and reduced spatial overlap. However, the D36K mutation had an intermediate effect, preserving the overall helical structure while increasing conformational flexibility. These observations underscore the context-dependent impact of electrostatic perturbations on helix stability, with residues crucial for maintaining helix integrity.

Furthermore, our simulations revealed that the conformational state of the TAP influenced the structural behavior of the luminal helix UL49.5. In the kinked TAP conformation, representing the minor cryo-EM population (27%), increased conformational heterogeneity was observed in all analyzed variants of the luminal helix, leading to reduced positional stability and partial shortening of helical segments. These findings suggest that the structural properties of the luminal helix are dependent on the conformational state of TAP, highlighting the dynamic nature of the UL49.5–TAP complex. The analysis of the UL49.5-TAP interaction interface emphasized the importance of specific polar residues within the UL49.5 *N*-terminal helix. However, our results also suggest that the salt bridges formed between RR(30–31) and D36K in UL49.5 and the TAP residues identified by MD simulations are unlikely to represent the only specific polar interactions within the UL49.5 *N*-terminal domain responsible for guiding the inhibitor into the TAP cavity. The highest number of strong polar interactions distributed evenly across the UL49.5 structure was observed in UL49.5-TAP complex in the unkinked conformation. Electrostatic contacts involving residues such as R6, R22, and R45 may play equally important roles. Reverse charge mutations in the luminal domain altered the interaction interface, affecting the stability and conformation of the UL49.5 within the complex, yet they created new, potentially compensating polar interactions, indicating the tendency of the interacting TAP regions to form this type of contact sites. However, these altered polar interaction networks included potentially compensatory salt bridges that may preserve the positioning of the N-terminal luminal domain within the ER-luminal TAP channel. Such preserved positioning could maintain the ability of UL49.5 to block antigenic peptide translocation, despite local remodeling of the interaction interface. This interpretation is consistent with the cell-based validation showing that the tested mutations did not abolish UL49.5-mediated MHC class I downregulation.

Moreover, the analysis of the cytoplasmic *C*-terminal region of UL49.5 and the exposure of the degron provided insights into the conformational dynamics and accessibility of this critical segment to KLHDC3. The exposure of the degron was controlled by the conformational coupling between UL49.5 and TAP, modulated by mutations within the luminal region of UL49.5. The highest stability of the *N*-terminal helix was observed in the complex of wild-type UL49.5 with unkinked TAP, resulting from strong polar interactions between the two proteins. Noticeably, in this complex, the *C*-terminal segment of UL49.5 also remained stable throughout the entire MD simulation (Figure 8S). The salt bridges between the *C*-terminus of the UL49.5 and TAP further enhanced this stability, as reflected by lower RMSF and SASA values (Figure 8S and 10S). In the complexes of UL49.5 mutants, high structural flexibility led to increased SASA values for the *C*-terminus and this excessive mobility may hinder the degron recognition by KLHDC3. Most importantly, in the complexes with mutants, the TM helix, preceding the *C*-terminus was located closer to TAP. The close proximity of the UL49.5 *C*-terminus to the TAP NBD domain may create steric hindrance making it impossible for KLHDC3 to approach the degron thereby preventing the recruitment of the CUL2 E3 ligase. Consistent with this model, the biological experiments confirmed that N-terminal electrostatic interactions in UL49.5 regulate TAP degradation. The two *N*-terminal mutants, UL49.5-D36K and UL49.5-RR(30–31)DD, were designed to disrupt local charge interactions and test their functional consequences. Immunoblotting and flow cytometry showed that these mutations abolished UL49.5-mediated TAP degradation while preserving the ability of UL49.5 to downregulate MHC class I surface expression. These findings indicate that TAP degradation and MHC class I downregulation can be functionally uncoupled and support an allosteric role of the *N*-terminal luminal region in controlling *C*-terminal degron-dependent TAP degradation. The biological experiments conducted in this study confirmed the importance of *N*-terminal salt bridges in UL49.5 in regulating TAP degradation. Mutations disrupting these electrostatic interactions which significantly allosterically changed the structural dynamics of the UL49.5, especially in the TM helix and the *C*-terminal region, prevented UL49.5-driven TAP degradation, underscoring the significance of these interactions in viral immune evasion mechanisms.

In conclusion, our study defines the structural and functional role of specific electrostatic determinants within the luminal domain of the UL49.5 protein in the inhibition of TAP and downregulation of MHC class I loading. The findings contribute to our understanding of the complex interplay between UL49.5 and TAP in viral immune evasion and highlight the importance of specific structural motifs for their interactions. Our results indicate that mutations introduced in the *N*-terminal region of UL49.5 can significantly impact the structure of the entire protein. Consequently, the closer positioning of the TM helix to TAP together with the high mobility of the *C*-terminus hinders KLHDC3 access to the degron, thereby preventing E3 ligase-mediated TAP degradation. Among the complexes with wild-type UL49.5, that with unkinked TAP appears to be the preferred, lowest-energy conformation characterized by a stable N-terminal helix, a correctly positioned TM helix, and a moderately flexible *C*-terminus, which can be easily recognized by KLHDC3.

More research is needed to explore the precise molecular mechanisms by which these electrostatic interactions regulate MHC class I downregulation and TAP degradation, with potential implications for the development of novel therapeutic strategies targeting viral immune evasion mechanisms.

## Acknowledgments

Computations were carried out using the computers of the Tricity Academic Supercomputer & Network Center of Informatics.

## Data Availability

NMR data have been deposited in the Biological Magnetic Resonance Data Bank (BMRB) under accession number 53843 UL49.5-D36K and 53844 UL49.5- RR(30–31)DD

## CRediT author statement

<u>Natalia Karska:</u> Supervision, Conceptualization, Methodology, Investigation, Formal analysis, Visualization, Writing – original draft, Writing – review and editing, Resources, Project administration, Funding acquisition.

<u>Małgorzata Graul:</u> Methodology, Investigation, Formal analysis, Validation.

<u>Igor Zhokov:</u> Methodology, Investigation, Formal analysis, Data curation, Writing – review and editing.

<u>Sylwia Rodziewicz-Motowidło</u>: Resources, Conceptualization, Writing – review and editing.

<u>Andrea D. Lipińska:</u> Conceptualization, Methodology, Investigation, Formal analysis, Validation, Supervision, Resources, Writing – review and editing, Funding acquisition.

<u>Magdalena J. Ślusarz:</u> Conceptualization, Methodology, Software, Formal analysis, Investigation, Visualization, Writing – review and editing.

## 4. Experimental Section

### Solid-phase peptide synthesis

All peptides were synthesized using SPPS techniques using an automated microwave peptide synthesizer (Liberty Blue, CEM Corporation, Matthews, NC, USA). The peptides were synthesized in a ProTide resin (0.2 mmol/g) using Fmoc chemistry and standard amino acid derivatives (Table 1). **Peptide cleavage** from the resin was performed using a mixture of 88% trifluoroacetic acid (TFA), 5% phenol, 5% deionized H_2_O and 2% triisopropyl silane (TIPS) (10 ml for 1 g of resin). The reaction ran for 4 h at ambient temperature. After 4 h the solution was filtered from the resin, concentrated under vacuum and treated with ice-cold Et_2_O to precipitate the peptide. The precipitated peptides were centrifuged for 10 min at 4000 rpm at 4°C, the Et_2_O phase was decanted (this step was repeated three times). Crude peptides were dissolved in deionized H_2_O and lyophilized.

### Peptide purification and characterization

The crude peptides were purified using the RP-HPLC technique on a semi-preparative Jupiter C18 column (250 mm × 10 mm, 5 µm) column from Phenomenex (Torrance, CA, USA). The mobile phase used during peptide purification consisted of (A) 0.1% trifluoroacetic acid (TFA) in deionized H_2_O and (B) 80% acetonitrile (ACN) in deionized H_2_O containing 0.1% trifluoroacetic acid (TFA) (Table 2). The flow rate was 5 ml/min. The separation process was monitored by UV absorbance at 223 nm. The purity of all the peptides was higher than 99% as estimated by RP-HPLC.

The purity of the peptide was confirmed by (i) RP-HPLC using a Kromasil C8 analytical column (250 mm × 4.6 mm, 5 µm) analytical column, using a linear gradient of 5 % B to 100% B over 60 min, where (A) was 0.1% (v:v) TFA in deionized H_2_O and (B) was 80% ACN in H_2_O containing 0.1% (v:v) TFA, and (ii) the MALDI-TOF MS spectra were acquired on a Bruker Biflex III MALDI-TOF spectrometer using an ultraviolet laser source (nitrogen, 337 nm); α-cyano-4-hydroxycinnamic acid was used as the matrix.

### CD measurements

CD studies were performed in DPC micelles (5 mM and/or 100 mM, POCH, Poland) in water (pH 7.0) or PBS (pH 7.4). The peptide solutions in the DPC micelles were prepared according to standard procedures. The CD spectra were recorded for each peptide (0.15 mg/mL) in the range of 185–260 nm with a Jasco J-815 spectropolarimeter (Jasco, Easton, USA). For each peptide, the CD spectra were recorded three times, at 30 °C using a 1-mm cell and are shown as the mean molar ellipticity of the residue (MRME, degree×cm2×dmol−1) versus wavelength λ (nm).

### Multidimensional NMR spectroscopy

The typical NMR samples were prepared by dissolving around 1.0 mM of peptide in a solution containing 100 mM of DPC-d38 dissolved in 90%/10% H_2_O/D_2_O. Multidimensional NMR data were acquired on Varian Inova 500 (1H frequency 500.606 MHz) NMR spectrometer (Palo Alto, California, USA), equipped with three channels, Performa IV z-gradient unit, and ^1^H/^13^C/^15^N probe head with inverse detection. Assignments of ^1^H, ^13^C, and ^15^N resonances were achieved through joint inspection of the 2D homonuclear TOCSY, collected with 80 ms, and NOESY, conducted with 150 ms – NMR experiments. These data were supplemented with 2D heteronuclear ^1^H-^13^C and ^1^H-^15^N HSQC spectra acquired on natural abundance of ^13^C and ^15^N isotopes. The ^1^H, ^13^C, and ^15^N resonances were referenced indirectly to external sodium 2,2-dimethyl-2-silapentane-5-sulfonate (DSS) using the 0.251449530 and 0.101329118 ratios for ^13^C and ^15^N, respectively. All collected NMR data were processed with NMRPipe software (13) and analyzed with the NMRFAM-SPARKY software (14). The assignment procedure resulted in more than 85% of ^1^H, ^13^C, and ^15^N resonances presented in all analyzed peptides were defined for specific residues.

### Evaluation of high-resolution 3D structure of the UL49.5 protein in solution

Initial structure calculations of the mutants UL49.5^22-56^RR(30–31)DD and UL49.5^22-56^D36K were performed with the CYANA software suite (version 3.98.15) (15,16). Analysis of the NOESY data recorded with a mixing time of 150 ms yielded 1017, 451, and 467 cross-peaks for the mutants UL49.5^22-56^RR(30–31)DD and UL49.5^22-56^D36K, respectively. Automatic NOEY assignment procedure (17) was performed together with conversion of NOE intensities into distance constraints provided to unique 267 (122 intra-residue, 71 sequential, 59 medium range and 15 long-range), 141 (80 intra-residue, 54 sequential, and 7 medium range), 128 (76 intra-residue, 40 sequential, and 12 medium range) upper limit distance constraints for all two mutants. These data were supplemented with 96 restrictions (UL49.5^22-56^RR(30–31)DD) and 65 restrictions (UL49.5^22-56^D36K) for the torsion angles of the backbone φ and ψ evaluated with TALOSn software (18) in chemical shifts assigned to 1H, 13C, and 15N base (Table S1). The distance constraints for hydrogen bonds were introduced on based geometrical criteria and defined as rHN-O = 2.1 ± 0.6 Å and rN-O = 3.0 ± 0.5 Å. The ensemble of 20 structures characterized by minimal residual target function were refined in a water shell with the YASARA program (19,20). The quality of the ensemble structures was analyzed with PROCHECK-NMR software (21) and WhatIf (22) (Table S1).

### All-atom MD simulation of UL49.5 in the membrane model

The three-dimensional structures of the TAP-UL49.5 complexes were retrieved from the Protein Data Bank (PDB codes: 9O94 and 9O9D) (12). The missing loop fragments and hydrogen atoms were added to the LEaP module of AMBER (23). The obtained structures of the TAP-UL49.5 complexes were copied and mutations were introduced to UL49.5 using AmberTools21 to generate six different complexes: TAP-UL49.5, TAP-UL49.5 UL49.5-RR(30–31)DD and TAP-UL49.5-D36K, each in both the kinked (9O94) and the unkinked (9O9D) conformation. In the next step, all complexes were embedded in the membrane model using the AmberTools PACKMOL-Memgen workflow and the Lipid21 force field (24,25). The lipid bilayer consisted of 200 lipid molecules, DMPC:DMPE:DMPS:POPC:POPE:POPS in a ratio of 27:15:8:27:15:8, to reproduce the conditions of the endoplasmic reticulum bilayer (26). The systems were hydrated with OPC water molecules (27) and the total charge of each system was neutralized with K^+^ counterions. The initial simulation boxes were 118 × 118 × 155 Å and 57256 water molecules, and 112 x 112 x160 Å and 59188 water molecules, for kinked/unkinked conformations, respectively. All systems were energy minimized and then subjected to molecular dynamics (MD) simulation. The system was first heated from 0 to 300 K with positional restrictions placed on the backbone atoms (weight of 5.0 kcal/mol /Å) followed by an equilibration for 5 ns. Finally, for each complex, 1 μs of the unrestrained MD was performed in AMBER 24 using the ff19SB force field (28). The long-range electrostatic interactions were computed using particle-mesh Ewald (PME) summation (29). The SHAKE algorithm was used to constrain covalent bonds involving hydrogen atoms (30). The temperature was regulated using a Langevin thermostat set at a target temperature of 300 K and a collision frequency of γ = 1.0 ps-1 (31). The pressure was regulated with a Berendsen barostat with an anisotropic pressure scaling for constant pressure (1 bar), and computations were carried out using the computers of the Center of Informatics Tricity Academic Supercomputer & Network.

The resulting trajectories were processed using the cpptraj module from the AMBER 24 package (32). The TAP-UL49.5 contact matrices were calculated in cpptraj using the native contact function with a distance cutoff of 4.0Å. Hydrogen bond analysis was performed in cpptraj using distance and angle criteria. A hydrogen bond was identified if the distance between the donor (X) and acceptor (Y) was not greater than 4.0Å and the XHY angle did not deviate from 135° more than ± 30°. The Root Mean Square Deviation (RMSD) and Root Mean Square Fluctuation (RMSF) were calculated using rms and the atomic function in cpptraj. The secondary structure content was calculated using the DSSP method of Kabsch and Sander (33). To estimate the Solvent Accessible Surface Area (SASA), the Linear Combination of Pairwise Overlaps (LCPO) method was used (34). The linear interaction energy (LIE) between UL49.5 and TAP was calculated using cpptraj. For van der Waals interactions, the default 12 Å cutoff was applied and the standard Lennard-Jones potential was used. The graphs were drawn using Python Matplotlib and Gnuplot 6.0.3 (35).

### Cell lines

The human melanoma Mel JuSo (MJS, a kind gift of Emmanuel Wiertz, UMCU, the Netherlands) and Madin-Darby bovine kidney (MDBK) cell line (American Type Culture Collection) were maintained in RPMI-1640 medium (Sigma-Aldrich), supplemented with 10% heat-inactivated fetal bovine serum, 2 mM L-glutamine, 100 IU/ml penicillin, 100 µg/ml streptomycin and 0.25 µg/ml amphotericin B (all from Thermo Scientific-Invitrogen), at 37 °C in 5% CO_2_ atmosphere.

### Antibodies

Rabbit polyclonal antibodies against a synthetic peptide derived from the *C*-terminal domain (H19) of UL49.5 were used to detect the UL49.5 protein (7). Cellular proteins were detected with the following antibodies: rabbit pAb anti-TAP1 (Proteintech), rabbit pAb anti-TAP2 (LSBio, USA), rabbit pAb anti-β-actin, (Novus), and mouse anti-human MHC class I mAb W6/32 (Santa Cruz Biotechnology).

### Generation of recombinant cell lines expressing variants of UL49.5

UL49.5 variants with point mutations were generated in the plasmid pcDNA3-IRES-NLS-GFP-UL49.5 (5) using the QuikChange Lightning Site-Directed Mutagenesis Kit (Agilent Technologies) according to the manufacturer’s protocol. The primers introducing mutations are listed in Table 2. The sequences were verified by DNA sequencing. Next, UL49.5 variants were amplified from the pCDNA3-IRES-NLS-GFP vector with the use of Q5® High-Fidelity DNA Polymerase (New England Biolabs) and primers 495BamFor and 495EcoRev (Table 4), and subsequently cloned in the *BamH*I-*EcoR*I restriction sites of the LZRS-IRES-GFP retrovirus vector (5). MJS cell lines were obtained by transduction with pantropic retroviruses generated in the GP2-293 cell line (Takara-Clontech), co-transfected with pLZRS plasmids and pVSV-G (Cell Biosystems) using TransIT-VirusGEN® Transfection Reagent according to the manufacturer’s instructions. Transduction was performed in the presence of 8 µg/ml polybrene (Merck) as previously described (7). GFP-positive cells were sorted using the FACSCalibur flow cytometer Sorting Option (Becton Dickinson). The following stable cell lines were constructed: MJS UL49.5 RR(30–31)DD and MJS D36K.

### Flow cytometry

Cell surface levels of MHC class I were examined by indirect staining with primary antibodies as indicated and, as a second step, goat anti-mouse IgG conjugated with phycoerythrin (PE) (Becton Dickinson). Cells were analyzed using the FACSCalibur flow cytometer (Becton Dickinson) and CellQuestPro software.

### Immunoblotting

Cells were resuspended in a lysis buffer containing 0.5% NP40 in 50 mM Tris-HCl pH 7.5, 5 mM MgCl_2,_, and the Complete mini protease inhibitor cocktail (Roche). Next, cell lysates were analyzed by SDS-PAGE and immunoblotting as previously described (7). The PVDF membranes were blocked overnight with 5% (w/v) skimmed milk in TBST (10 mM Tris-HCl, pH 8.0, 150 mM NaCl, 0.1% Tween 20) and incubated with suitable primary antibodies followed by extensive washing and incubation with secondary horseradish peroxidase-conjugated goat IgG (Jackson ImmunoResearch, USA). Proteins were detected using ECL (SuperSignal West Pico Plus substrate, Thermo Scientific or SignalBright Max Chemiluminescent Substrate, Proteintech). Densitometry was performed using UVIBand software integrated with the Alliance chemiluminescence analysis device (UVITEC, United Kingdom).

### Statistical analysis

Statistical analysis was performed by one-way ANOVA using Prism version 9.3.1 (GraphPad Software, USA).

