## Supplementary Figures for "Unlocking viral evasion: Luminal charge interactions in BoHV-1 UL49.5 allosterically control TAP degradation"

### Supplementary Materials

**A)**

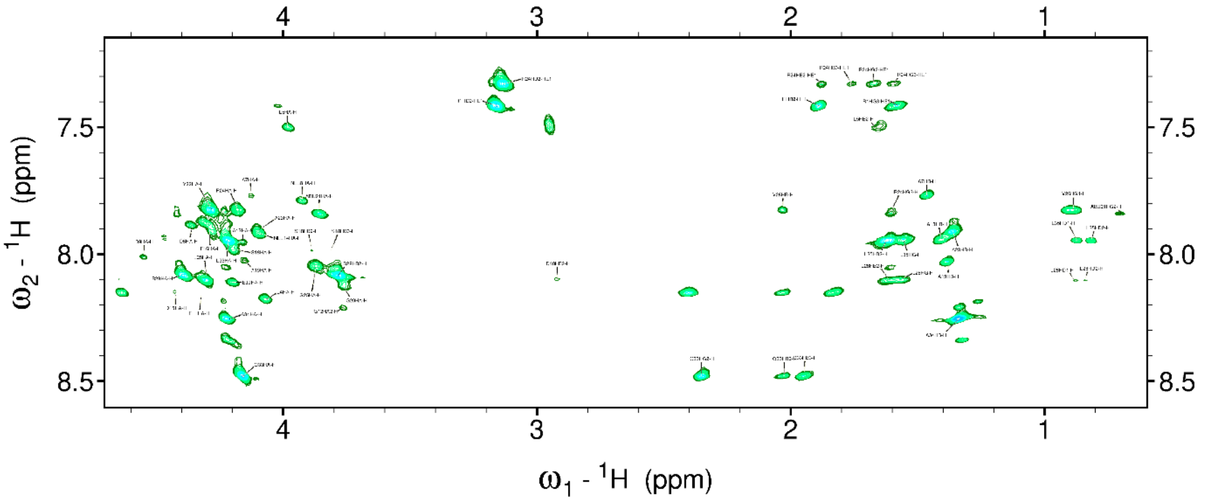


**B)**

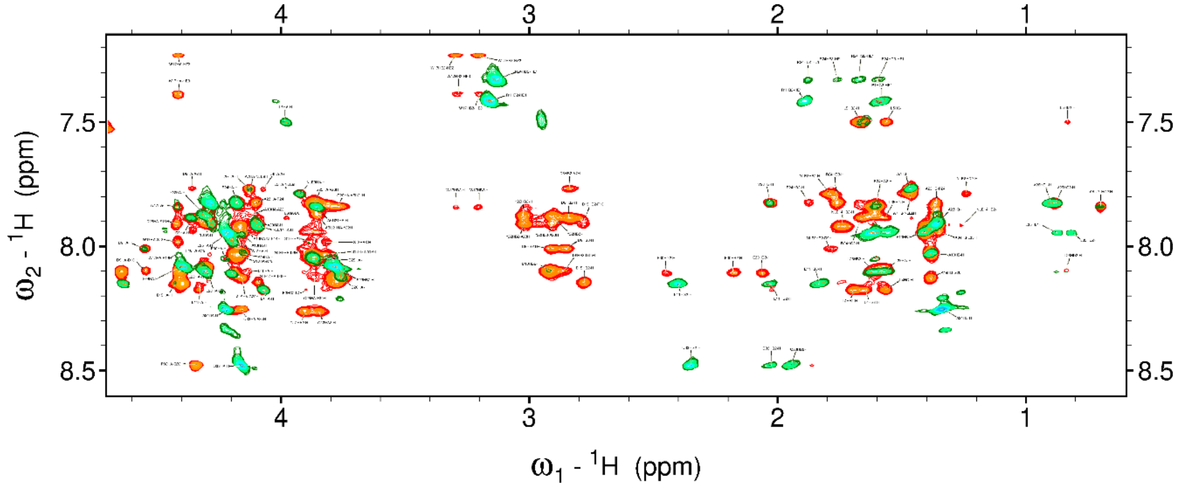


**Figure 1S**. 2D NMR spectra of the UL49.5$^{22\text{-}56}$RR(30–31)DD peptide in DPC-$_{d38}$ micelles (1.0 mM peptide in 100 mM DPC-$_{d38}$, 90%/10% - $H_2O/D_2O$) recorded at 30 °C. (**A**) Fingerprint region of the 2D TOCSY spectrum ($\tau_{mix}$ = 80 ms). (**B**) Fingerprint region of the overlaid 2D TOCSY (green, $\tau_{mix}$ = 80 ms) and NOESY (orange, $\tau_{mix}$ = 150 ms) spectra.

A)

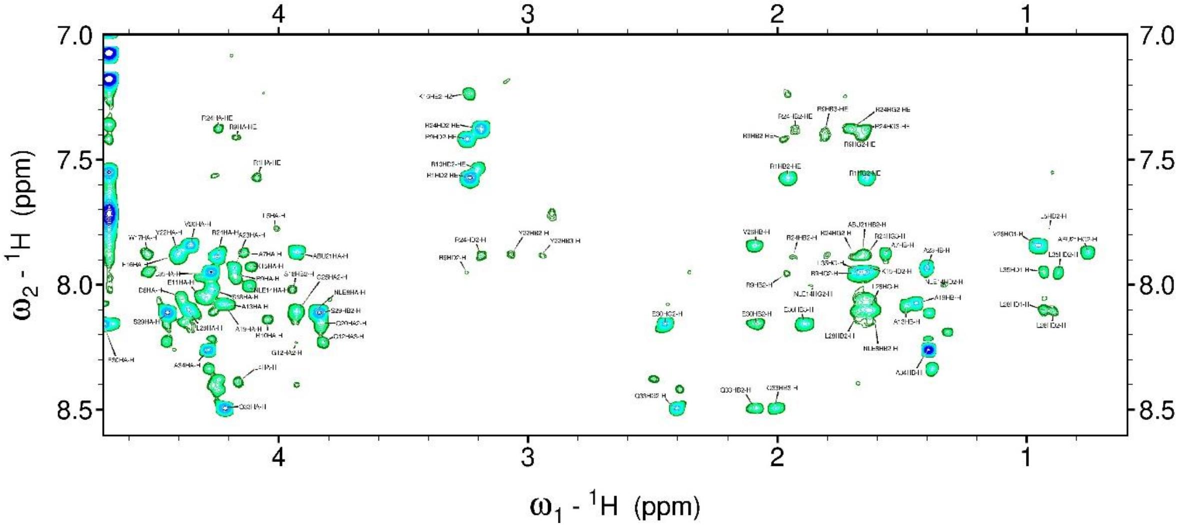


B)

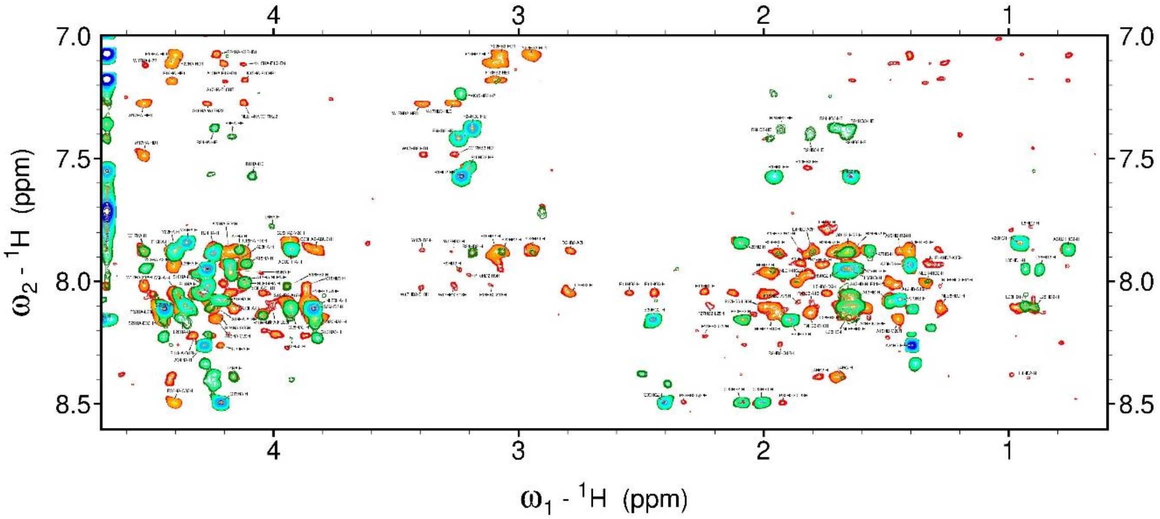


**Figure 2S.** 2D NMR spectra of the UL49.5$^{22\text{-}56}$D36K peptide in DPC$_{\text{-d38}}$ micelles (1.0 mM peptide in 100 mM DPC$_{\text{-d38}}$, 90%/10% - $H_2O/D_2O$) recorded at 30 °C. (**A**) Fingerprint region of the 2D TOCSY spectrum ($\tau_{mix}$ = 80 ms). (**B**) Fingerprint region of the overlaid 2D TOCSY (green, $\tau_{mix}$ = 80 ms) and NOESY (orange, $\tau_{mix}$ = 150 ms) spectra

A)

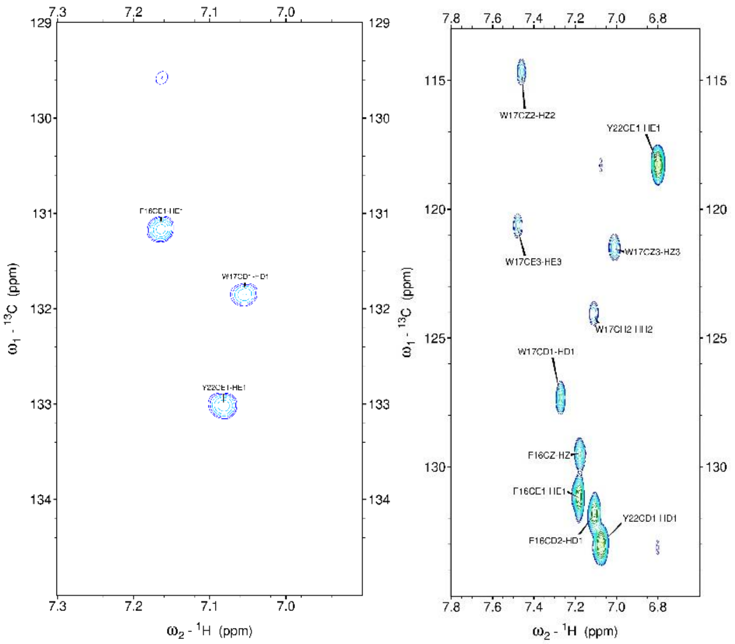


B)

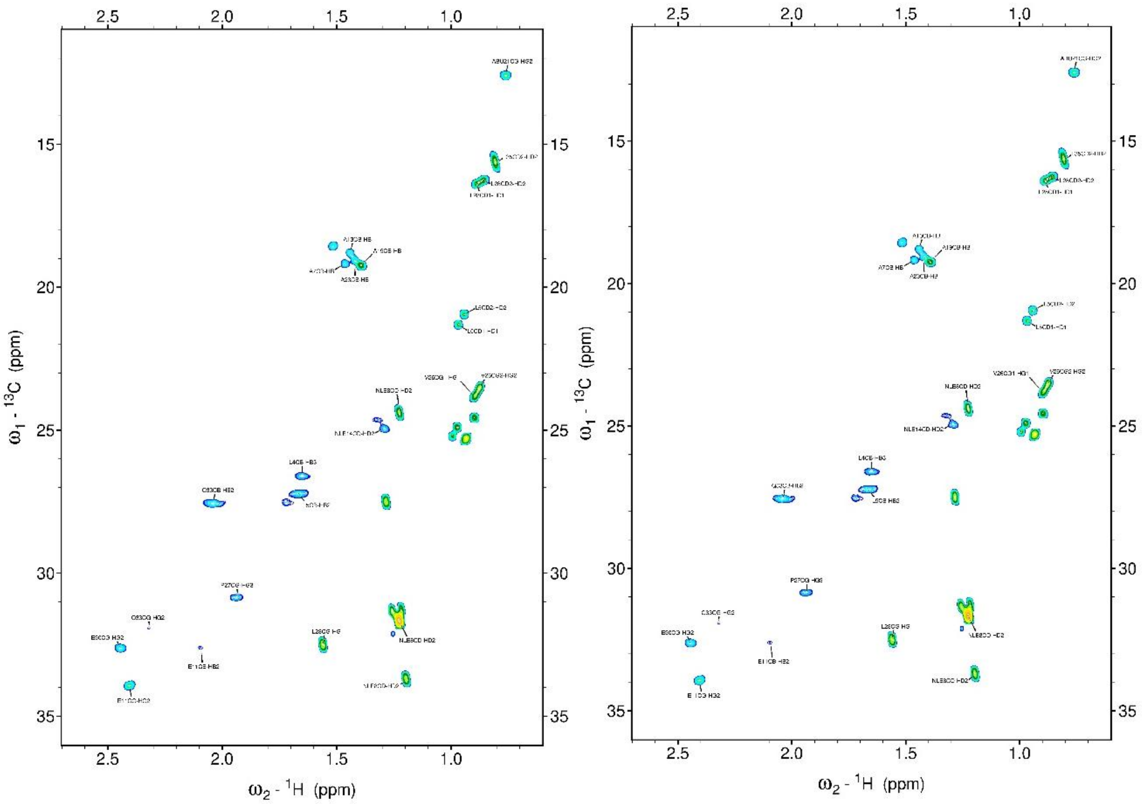


**Figure 3S.** 2D $^{1}H$–$^{13}C$ HSQC spectra of the UL49.5$^{22\text{-}56}$RR(30–31)DD peptide in DPC$_{\text{-d38}}$ micelles (1.0 mM peptide in 100 mM DPC$_{\text{-d38}}$ 90%/10% - $H_2O/D_2O$) recorded at 30 °C. (**A**) Fingerprint region of the aromatic side chains. (**B**) Fingerprint region of the aliphatic side chains.

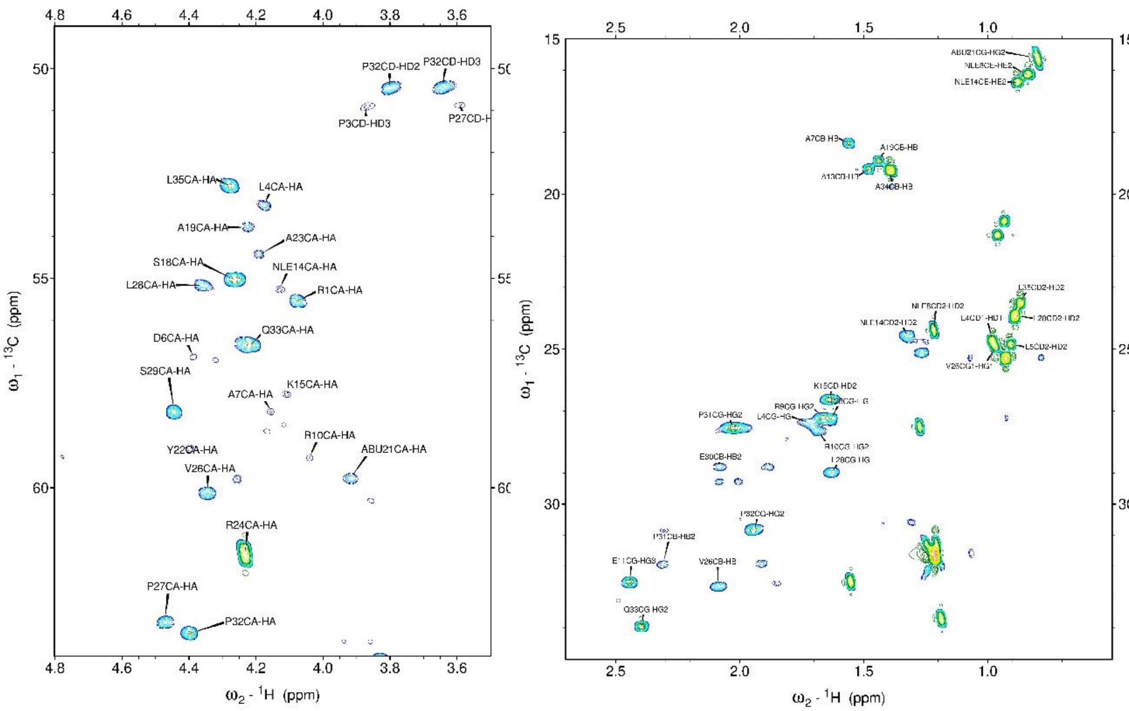

**Figure 4S.** 2D $^1H$–$^{13}C$ HSQC spectra of the UL49.5$^{22\text{-}56}$D36K peptide in DPC-$_{d38}$ micelles (1.0 mM peptide in 100 mM DPC-$_{d38}$, 90%/10% - $H_2O$/$D_2O$) recorded at 30 °C. Fingerprint region of the aliphatic side chains.

A)

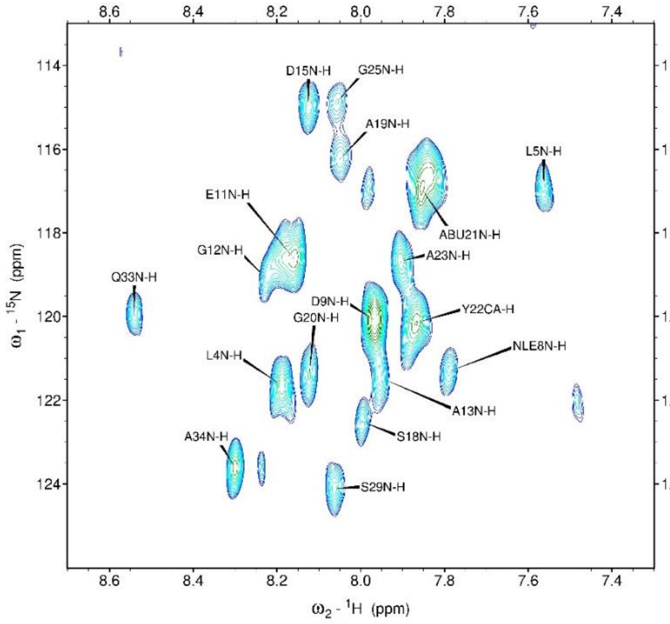


B)

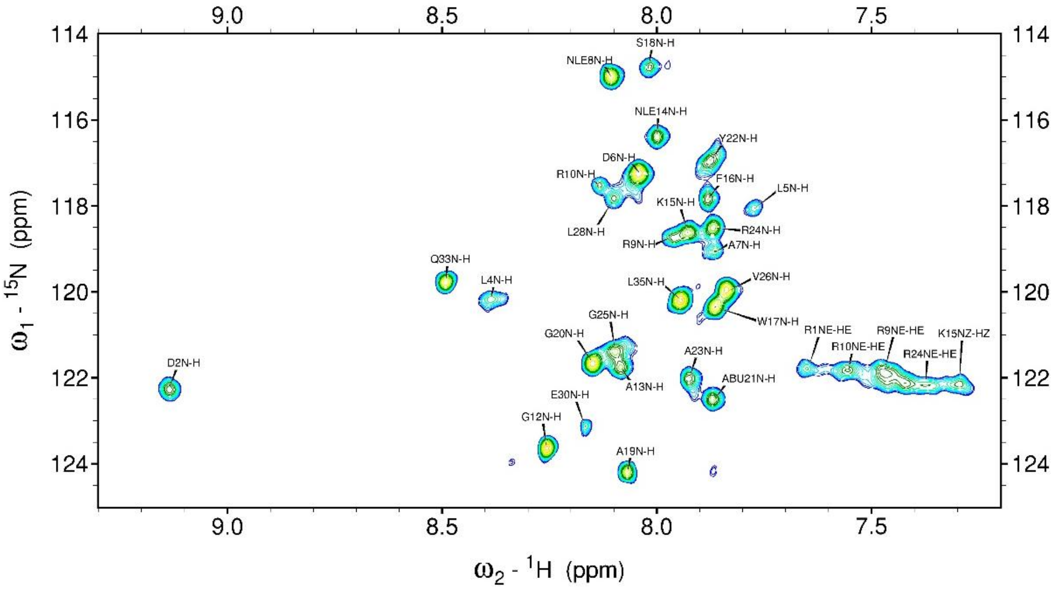


**Figure 5S.** Fingerprint regions of the 2D $^{1}H$–$^{15}N$ HSQC spectra of (A) UL49.5$^{22\text{-}56}$RR(30–31)DD and (B) UL49.5$^{22\text{-}56}$D36K in DPC$_{\text{-d38}}$ micelles (1.0 mM peptide, 100 mM DPC$_{\text{-d38}}$, 90%/10% - $H_2O/D_2O$) recorded at 30 °C.

A)

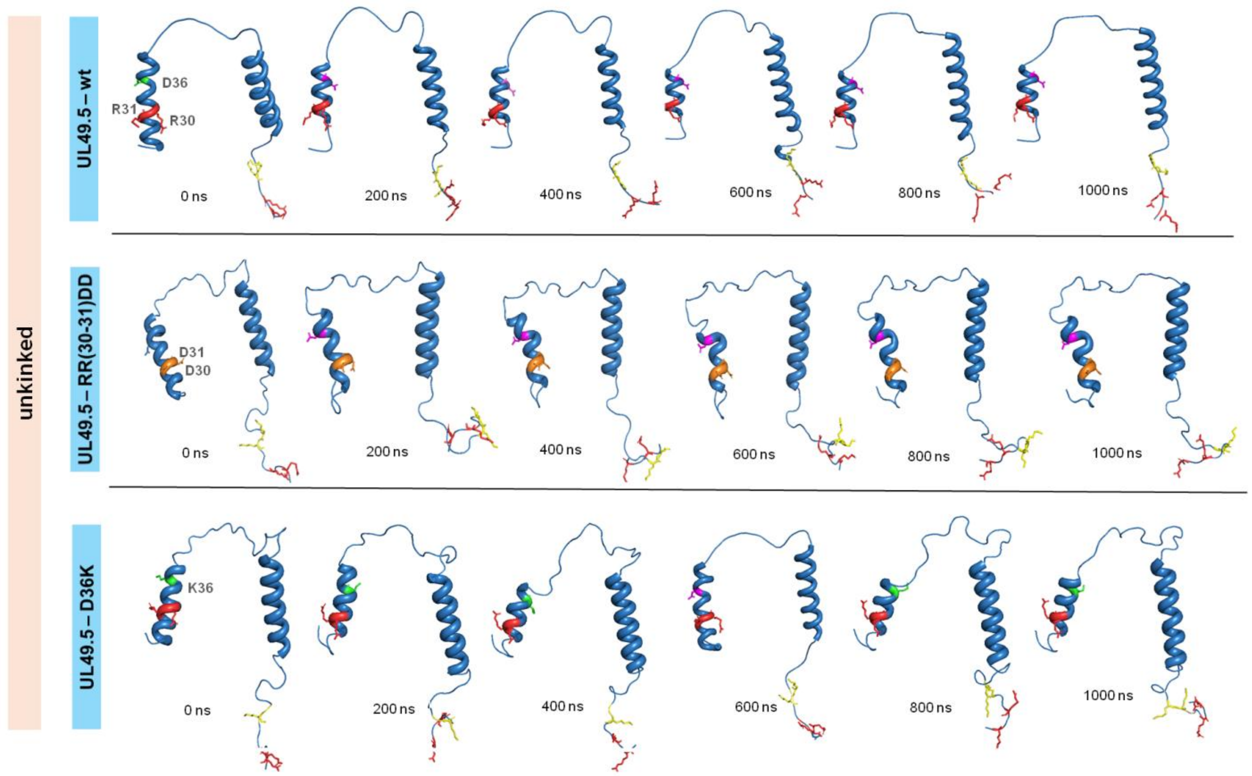


B)

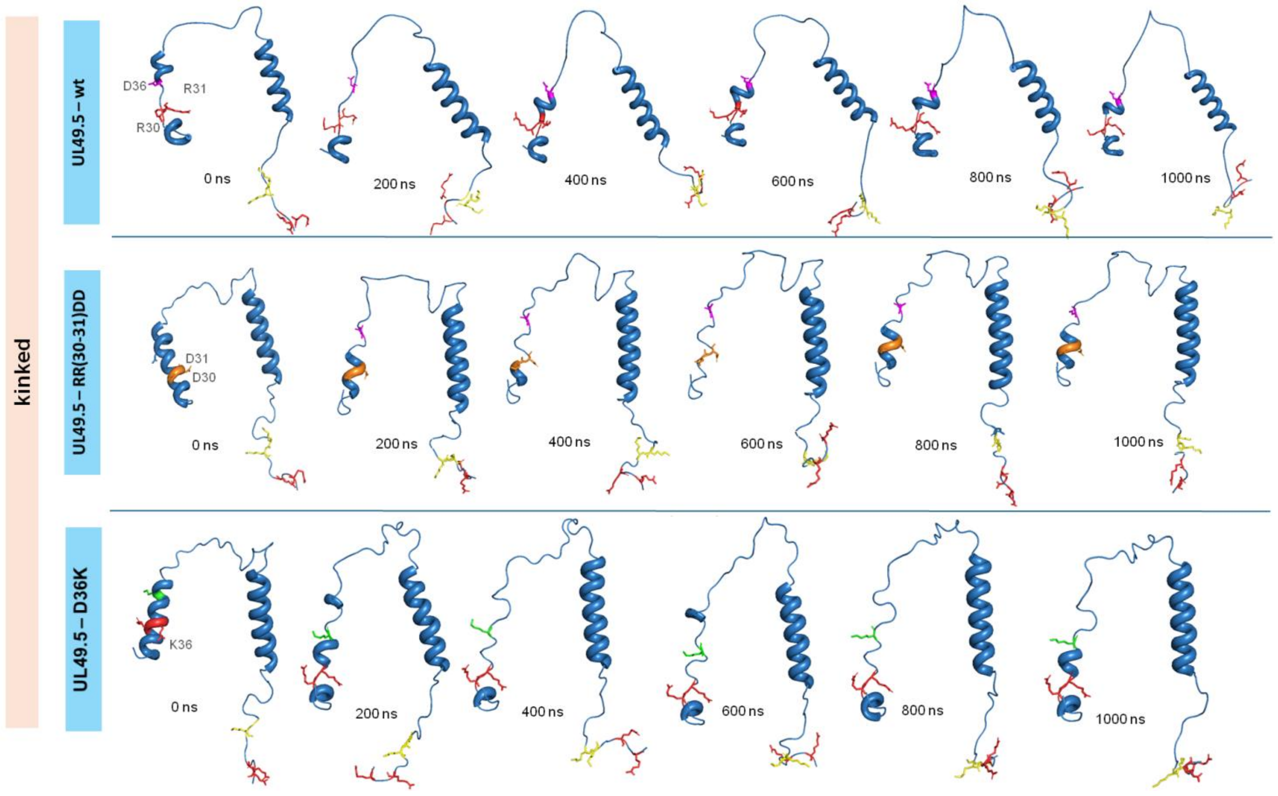


**Figure 6S. Time evolution of UL49.5 and its mutants in complex with TAP during 1 μs molecular dynamics simulations.** Representative snapshots at 0, 200, 400, 600, 800, and 1000 ns are shown for wild-type UL49.5 (top), UL49.5-RR(30–31)DD (middle), and UL49.5-D36K (bottom). Simulations were performed with TAP in the (A) unkinked and (B) kinked conformations. The protein backbone is shown as a blue cartoon, and selected residues are shown as sticks. In the unkinked conformation, wild-type UL49.5 maintains a stable luminal α-helical segments, whereas the

RR(30–31)DD and D36K mutants display altered helix continuity and increased conformational variability. In the kinked TAP conformation, all variants exhibit enhanced structural heterogeneity, particularly within the luminal region, accompanied by partial fragmentation of helical segments, while the transmembrane helix remains largely stable across all systems.

**A)**

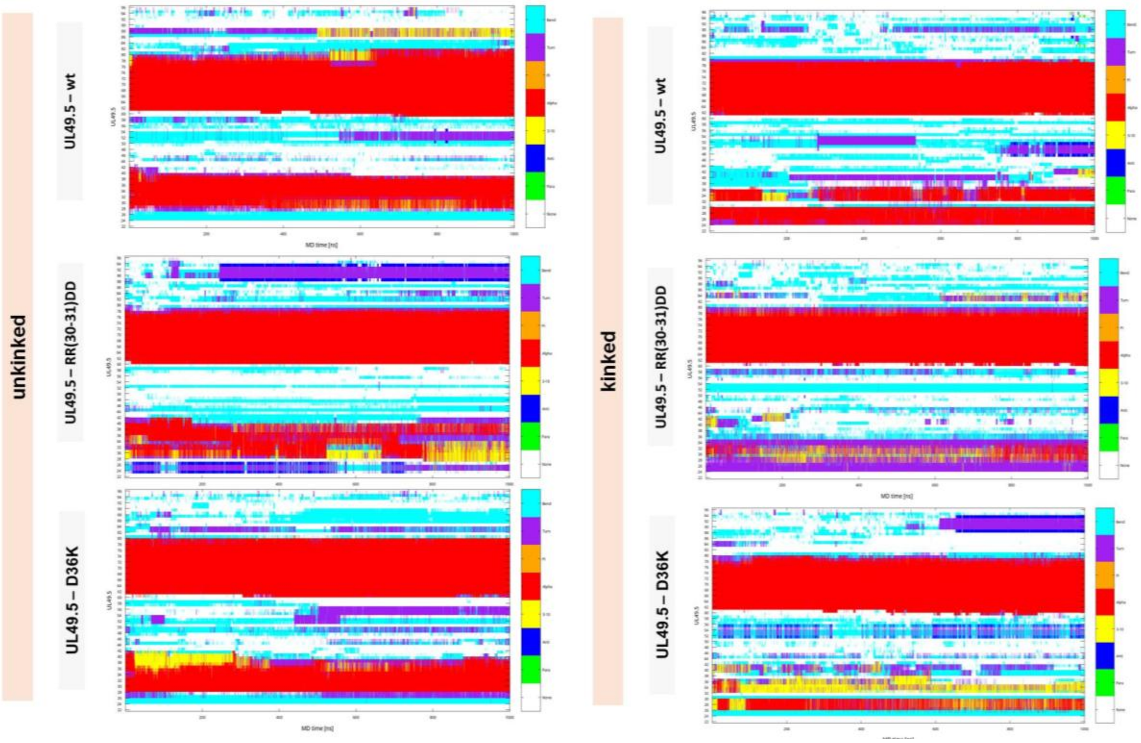


**B)**

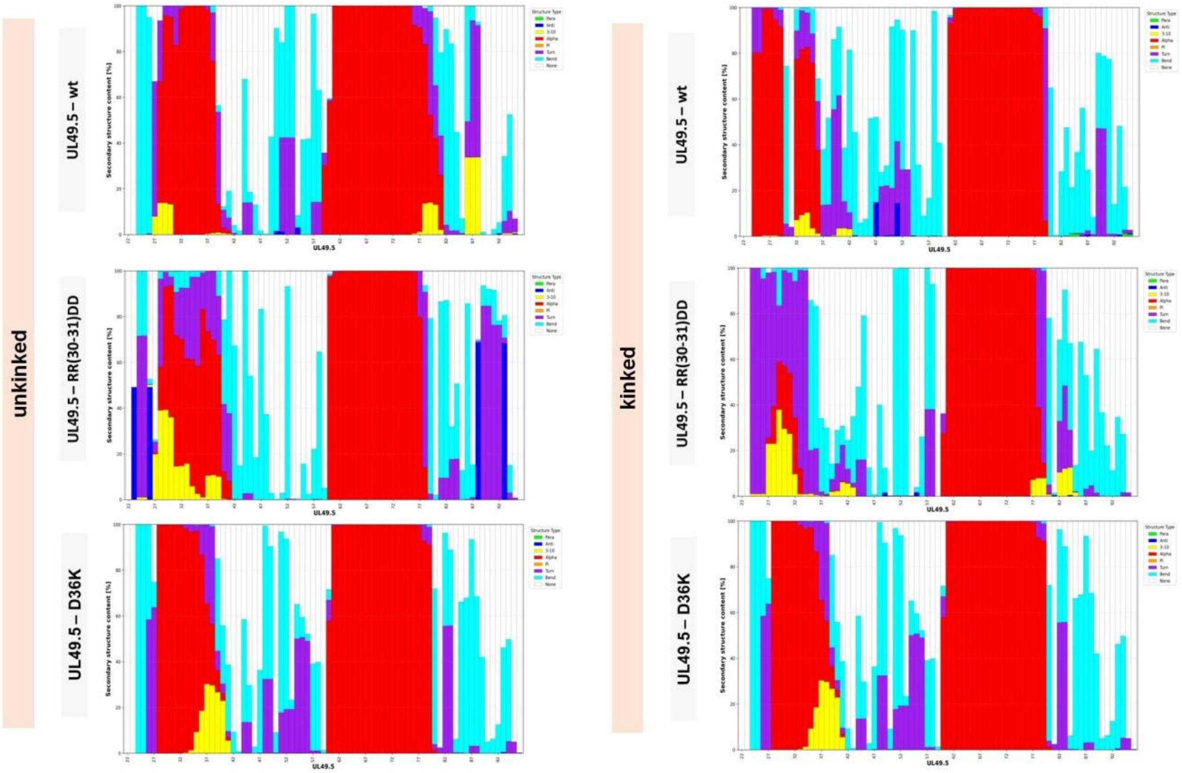


**Figure 7S. The secondary structure analysis (DSSP) of UL49.5-wt and its mutant variants in complexes with TAP in the unkinked and kinked conformation during 1 µs molecular dynamics simulations. (A)** Time-resolved DSSP secondary structure map for each residue over the simulation trajectory. **(B)** Per-residue secondary structure composition expressed as percentage occupancy based on DSSP analysis.

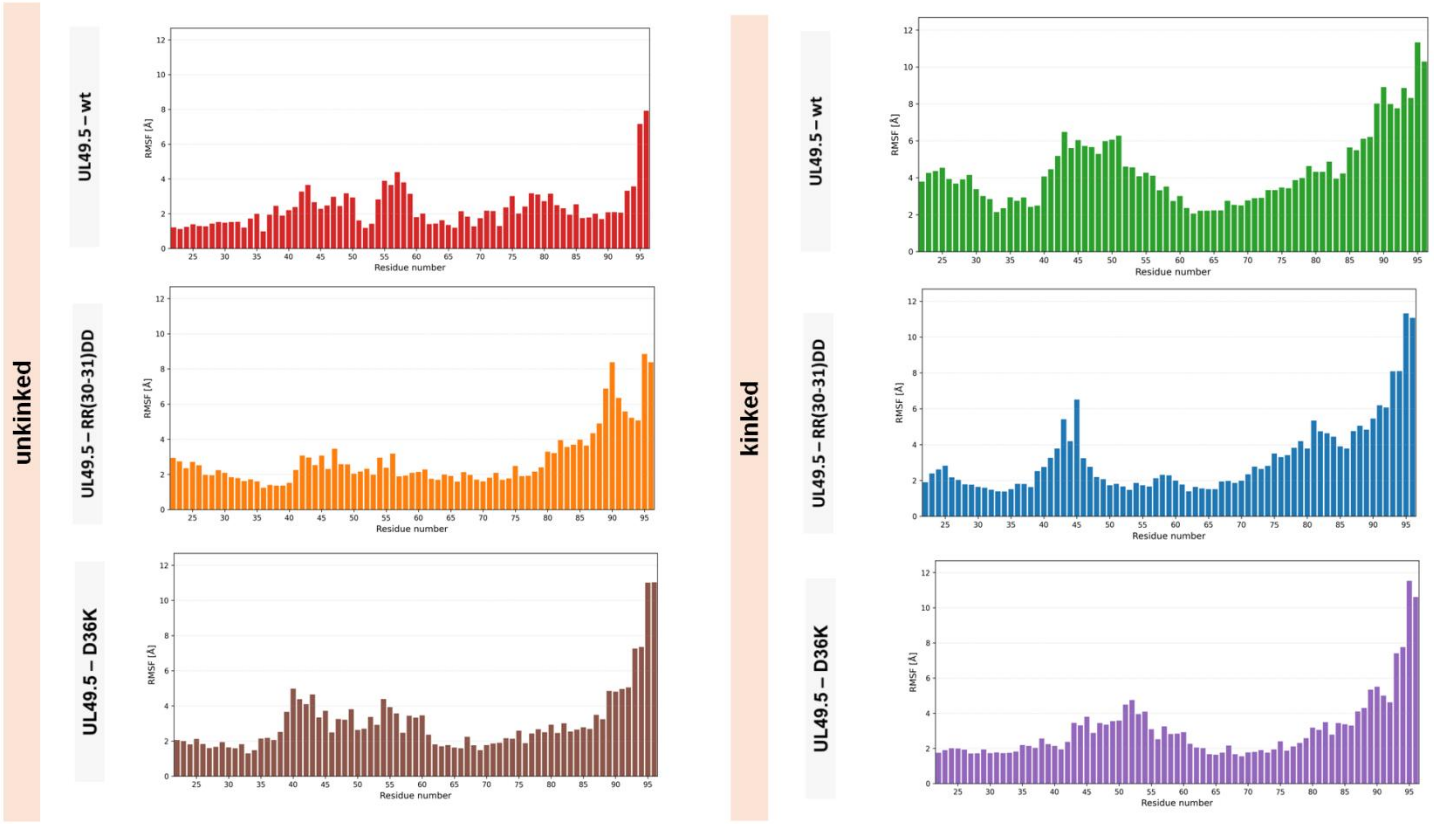


**Figure 8S. Per-residue root mean square fluctuation (RMSF) of UL49.5-wt and its mutant variants (RR(30–31)DD and D36K) in complex with TAP in the unkinked and kinked conformational states during 1 μs molecular dynamics simulations.** RMSF analysis of the wild-type UL49.5 protein in the unkinked TAP conformation indicates moderate fluctuations across both the luminal and transmembrane regions. Both, UL49.5-RR(30–31)DD and UL49.5-D36K mutants exhibit higher RMSF values than the wild-type protein, particularly within the cytoplasmic region, while within the luminal and transmembrane regions moderate fluctuations are observed. This behavior contrasts with the wild-type protein, in which the cytoplasmic region does not show such a clear increase in flexibility compared to the rest of the structure. In the kinked TAP conformation in all complexes the largest fluctuations occur within the cytoplasmic *C*-terminal segment. The remaining regions exhibit substantially lower RMSF values, consistent with the preserved α-helical organization of the transmembrane segment and the partially retained helical propensity within the luminal region

**A)**

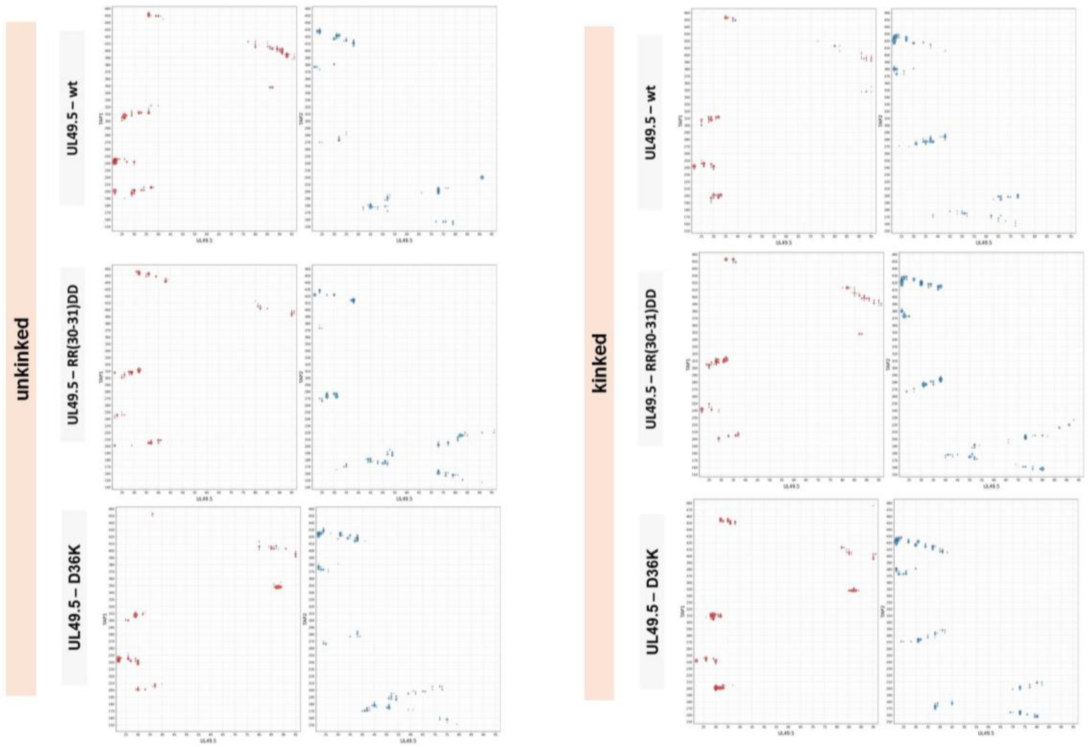


**B)**

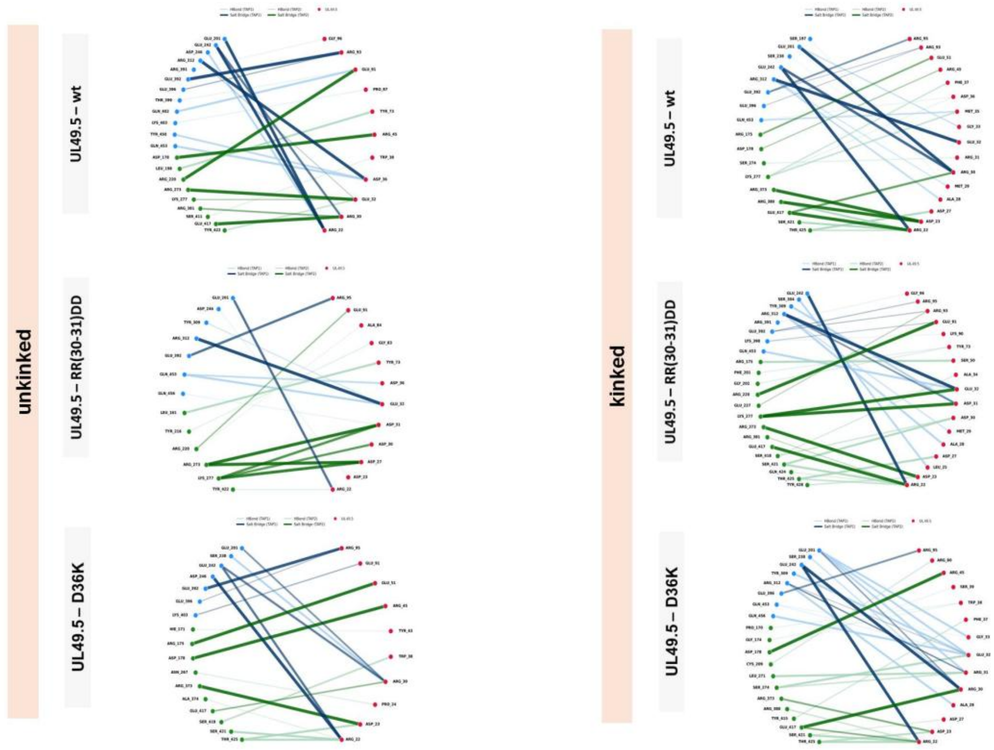


**Figure 9S. Hydrogen bond and salt bridge interaction networks between UL49.5 (wt and mutants) and TAP in the unkinked and kinked conformational states during 1 μs molecular dynamics simulations. (A)** All detected hydrogen bonds and salt bridges **(B)** The most persistent and strongest hydrogen bonds and salt bridges, selected based on their frequency of occurrence and stability. The highest number of strong salt bridges was observed in the wild-type UL49.5 complex with unkinked TAP (see Figure 10S). These salt bridges are evenly distributed across the UL49.5 structure, engaging both the N- and C-termini which results in enhanced overall stability of this complex.

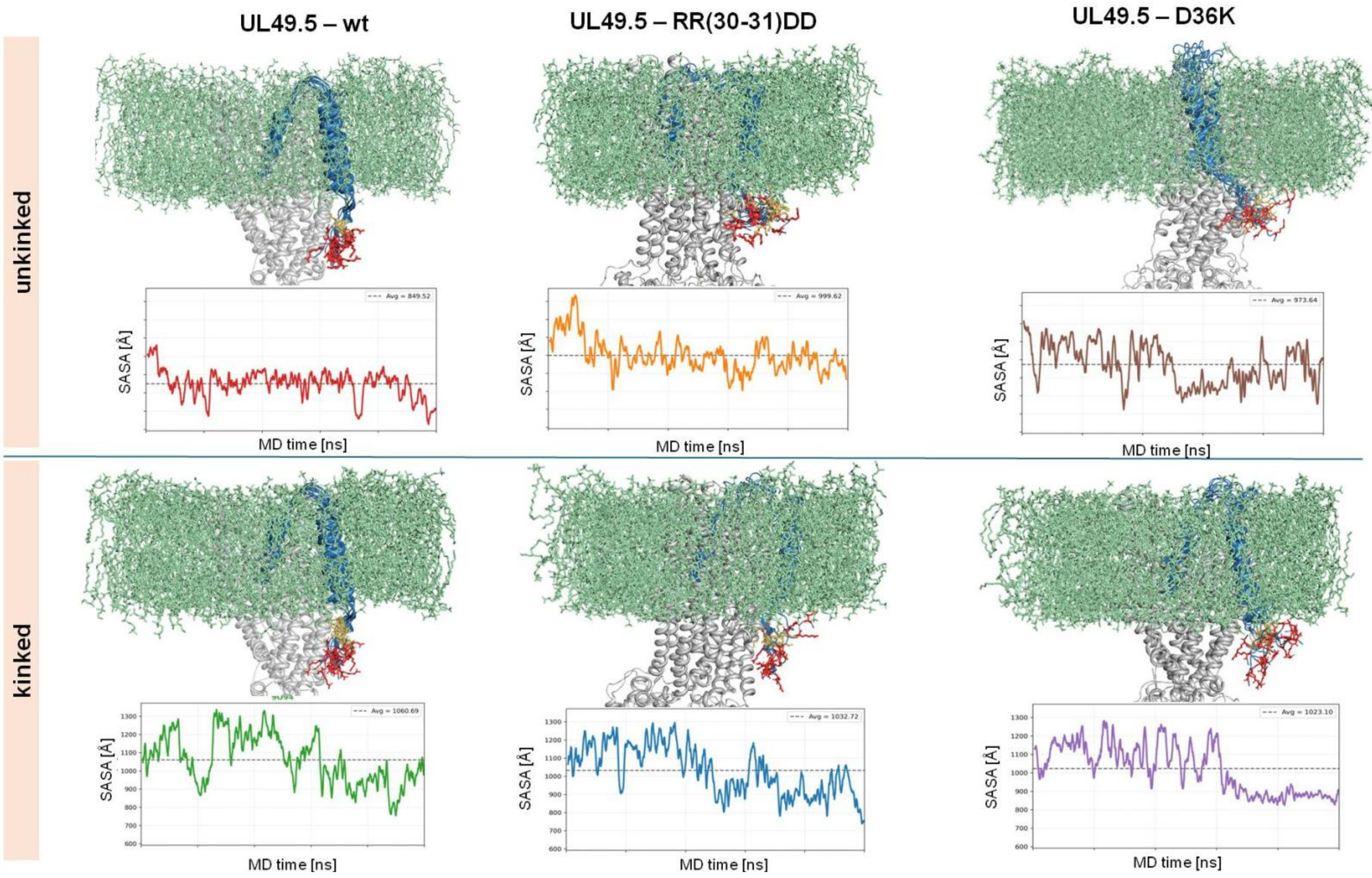


**Figure 10S. Conformational arrangement and solvent accessibility of the UL49.5 cytoplasmic *C*-terminal region in membrane-embedded UL49.5–TAP complexes.** UL49.5-wt, UL49.5-RR(30–31)DD and UL49.5-D36K are shown in complexes with TAP in the unkinked (top) and kinked (bottom) conformations. UL49.5 is represented as a blue cartoon, with residues R93 and R95 highlighted in red and residues K78 and K79 in yellow. The TAP transporter is shown in grey, and the lipid bilayer is depicted as green sticks. The lowest average SASA value, 849.52 Å², was recorded for the C-terminus of the wild-type UL49.5 complexed with unkinked TAP. This differs significantly from the values obtained for the other systems here high structural flexibility led to increased average SASA values.

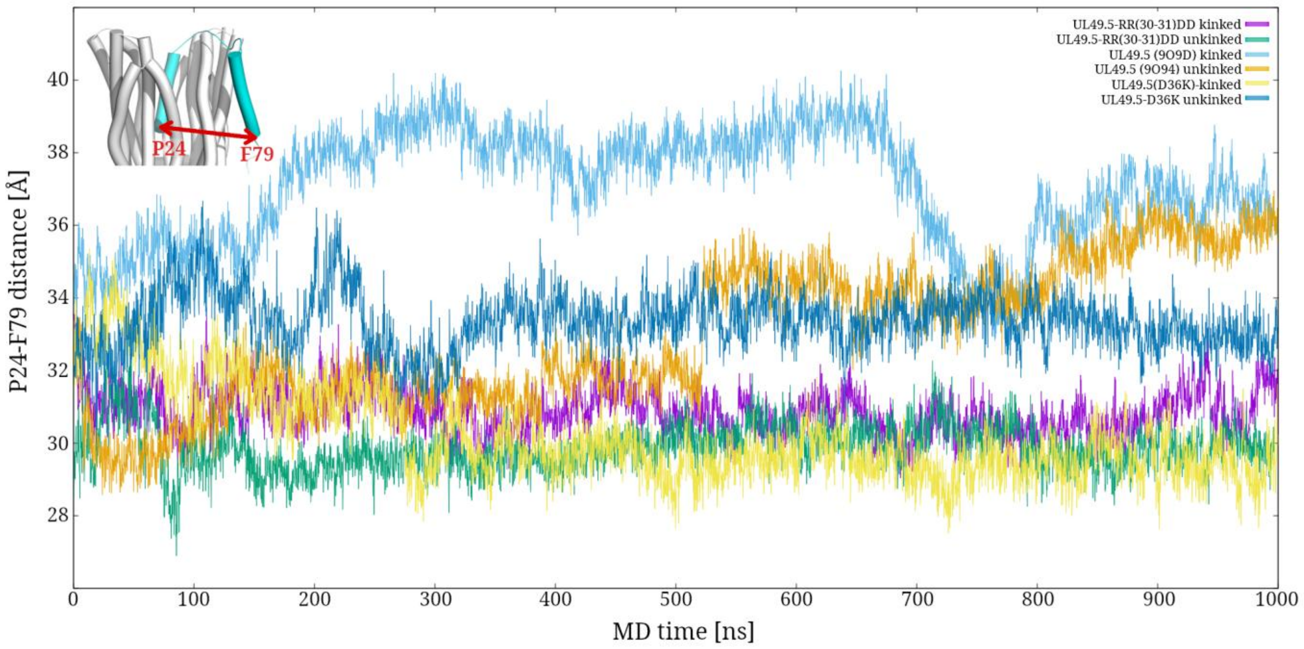


**Figure 11S. Distance between the termini of the UL49.5 helices (P24-F79). Left top corner: TAP is represented as light grey cylinders and UL49.5 as cyan cylinders. Red arrow indicates the measured distance.** The most significant fluctuations throughout MD simulation were observed for both complexes of wild-type UL49.5. After 800 ns, the position of the TM helix stabilized, maintaining a P24-F79 distance of approximately 37 Å.
